# Y RNA-derived fragments in a complex with YBX1 modulate PARP1 residency at DNA double strand breaks

**DOI:** 10.1101/2024.06.18.599493

**Authors:** Annabelle Shaw, Kamal Ajit, Monika Gullerova

**Affiliations:** Sir William Dunn School of Pathology, University of Oxford, South Parks Road, Oxford, OX1 3RE, United Kingdom

**Keywords:** Y RNA, YBX1, PARP1, double strand breaks, m5C, NSUN2, auto-ADP-ribosylation, bystander effect, exosomes

## Abstract

To protect genome integrity from pervasive threats of damage and prevent diseases like cancer, cells employ an integrated network of signalling pathways called the DNA damage response. These pathways involve both protein and RNA components which can act within the damaged cell or be transferred intercellularly to influence population-wide responses to damage. Here, we show that radioprotection can be conferred by damage-derived exosomes and is dependent on YBX1-packaged Y3-derived ysRNA. In recipient cells, ysRNA are methylated on cytosine by an RNA methyltransferase NSUN2, and bound by m5C reader, YBX1. YBX1/ysRNA localises at double strand break (DSB) sites to promote efficient DNA repair and cell survival through complex formation with PARP1. YBX1 modulates PARP1 auto-modification by facilitating ysRNA ADP-ribosylation, promoting increased PARP1 residency at DSBs. Our data highlight an unprecedented role for these under-studied species of small non-coding RNA, identifying them as a novel substrate for PARP1 mediated ADP-ribosylation and their function in DNA repair.

## Introduction

The human genome is under constant threat of damage from endogenous and exogenous sources, resulting in genome instability if improperly repaired. Cells therefore utilise a complex network of signalling pathways, collectively termed the DNA damage response (DDR), to detect and repair these lesions, thus ensuring genome integrity and preventing diseases such as cancer(Groelly et al., 2023; Jackson and Bartek, 2009). DNA double strand breaks (DSBs) represent the most cytotoxic lesions which can be induced by ionising radiation (IR) and are repaired via two major pathways: fast, but error-prone, non-homologous end-joining or high-fidelity homologous recombination(Jackson and Bartek, 2009). In addition to protein effectors, there is increasing evidence for major roles of RNA, particularly non-coding RNA (ncRNA), in DSB repair. These functions can be classified as in *cis* or in *trans* to DSBs(Bader et al., 2020).

It recently emerged that DDR can be communicated intercellularly via exosomes, a subset of extracellular vesicles (EVs) with a diameter of 40-150 nm(Zhang et al., 2015). Many studies have reported alterations to exosome cargo, including protein and RNA, following IR treatment of the cells from which they are secreted (Abramowicz et al., 2020; Abramowicz et al., 2019; Jelonek et al., 2015; Mutschelknaus et al., 2017; Yentrapalli et al., 2017). Results of such cargo transfer include protection of healthy bystander cells, which are in the vicinity of irradiated cells but are not exposed themselves, against subsequent DNA damaging insult(Mrowczynski et al., 2018; Mutschelknaus et al., 2016). Such intercellular communication generates a co-operative response to DNA damage, particularly recurring insults, which can increase the overall survival of a cell population(Shaw and Gullerova, 2021).

The multifunctional nucleic acid binding protein, Y-Box Binding Protein 1 (YBX1), is capable of binding to various forms of DNA and RNA(Hasegawa et al., 1991) and has been implicated in packaging of messenger RNA (mRNA), microRNA (miRNA) and other small non-coding RNA (sncRNA) species into EVs(Gopanenko et al., 2020; Liu et al., 2021; Shurtleff et al., 2016; Shurtleff et al., 2017). Among many other functions, YBX1 is also involved in DNA repair. While the full extent of YBX1 involvement in DNA repair is still under study, it is known to translocate to the nucleus upon genotoxic stress, associate with DNA damage markers such as Ser139-phosphorylated histone H2AX (γH2AX), and participate in DNA repair protein complexes(Ghanawi et al., 2021; Kim et al., 2013; Sorokin et al., 2005). YBX1 therefore represents a strong candidate for involvement in intercellular radioprotection, due to its dual role in RNA packaging and DNA repair.

One example of YBX1-packaged exosomal RNA is Y RNA(Shurtleff *et al*., 2017). Y RNA are a class of ncRNA that are transcribed by RNA polymerase III from four genes in the human genome, giving rise to four RNAs from distinct promoters: Y1, Y3, Y4 and Y5. Ranging from 84-113 nucleotides (nt) in length, these RNA form hairpin structures that facilitate their interaction with effector proteins, including Ro60 and La(Hendrick et al., 1981; Kowalski and Krude, 2015; Wolin and Steitz, 1983). Y RNA functions are overall poorly characterised, although they reportedly include regulation of Ro60 subcellular localisation, RNA processing and quality control, DNA replication and stress responses, including promotion of cell survival following UV irradiation(Boccitto and Wolin, 2019; Chen et al., 2003; Sim et al., 2009). Full length Y RNA may also be processed into shorter fragments of 24-32 nt, termed Y RNA-derived fragments (ysRNA). This cleavage is facilitated by RNase L and may occur in a caspase-dependent manner during apoptosis(Billmeier et al., 2022; Donovan et al., 2017; Nicolas et al., 2012). However, the functions of these fragments remain largely unknown.

NSUN2 and DNMT2 are two essential RNA methyltransferases involved in the modification of cytosine to form 5-methylcytosine (m5C) in RNA. NSUN2, a member of the NOP2/Sun RNA Methyltransferase family, plays a crucial role in catalysing this methylation process across various RNA species, including tRNA, mRNA, and non-coding RNAs. DNMT2, specifically methylates cytosine at the 38th position in tRNA Asp (GTC), forming m5C. The activity of DNMT2 is essential for maintaining tRNA stability and ensuring proper decoding during translation(Tuorto et al., 2012). Additionally, both NSUN2 and DNMT2 play significant roles in stress response and are involved in DNA damage response pathways, highlighting its multifaceted role in cellular protection and repair mechanisms(Chen et al., 2020). YBX1, is an m5C reader. It specifically binds to m5C-modified RNA, influencing the fate of these RNAs within the cell. YBX1’s ability to recognize and bind m5C-modified RNA underscores its role in gene expression regulation and the cellular response to DNA damage(Chen et al., 2024).

In line with its DNA repair role, YBX1 has recently been reported as an interactor and regulator of PARP1(Alemasova et al., 2018; Naumenko et al., 2020). Primarily known as a DNA single strand break repair factor, PARP1 has recently also been implicated in DSB repair(Chen et al., 2019b; Ray Chaudhuri and Nussenzweig, 2017). Following activation, PARP1 can modify both itself and other target proteins by ADP-ribosylation, resulting in a readable signal that is capable of recruiting DNA repair proteins to the lesion, as well as enabling chromatin remodelling events that enhance repair(Chen *et al*., 2019b; Ray Chaudhuri and Nussenzweig, 2017; Smith et al., 2019; Smith et al., 2023). Beyond its role in modifying proteins, PARP1 can add ADP-ribose polymers to RNA target molecules. PARP1 plays a significant role in RNA polyadenylation, export, and stability, as well as ribosome biogenesis. During thermal stress, PARP1 binds and modifies poly(A) polymerases (PAP), decreasing global polyadenylation, which impacts RNA stability and export, thereby reducing protein translation efficiency. Additionally, PARP1 is involved in mRNA export through interactions with the THO/TREX complex and other receptors. It influences mRNA stability by regulating pathways like nonsense-mediated decay (NMD) and stabilizing certain mRNA-binding proteins. Furthermore, PARP1 is critical in ribosome biogenesis, impacting nucleolar architecture, ribosomal RNA processing, and ribosome assembly(Eleazer and Fondufe-Mittendorf, 2021). However, the role of PARP1 in modification of RNA at DSBs and its relevance to repair is not known.

Here, we report that exosome-mediated radioprotection is dependent on YBX1-packaged Y3-derived ysRNA which enter the nucleus of bystander cells upon DNA damage. These ysRNA are methylated by NSUN2 and interact with YBX1 at DSB sites to promote efficient DNA repair and cell survival through complex formation with PARP1. YBX1 facilitates ADP-ribosylation of ysRNA, modulating PARP1 auto-modification and therefore promoting increased PARP1 residency at DSBs to enhance DNA repair.

## Results

### Exosomes derived from damaged cells mediate protective YBX1 dependent bystander effect

Previous studies in human cells have shown a radioprotective bystander phenotype that is transmitted via exosomes, resulting in increased recipient cell proliferation and survival upon radiation insult when treated with exosomes derived from damaged donor cells(Mrowczynski *et al*., 2018; Mutschelknaus *et al*., 2016). We sought to determine whether this phenotype was mediated by enhanced DNA repair, using exosome transfer experiments (Figure 1A). Exosomes were isolated by size exclusion chromatography (Figure S1A-F) from the culture medium of HEK293T cells, which were either untreated or treated with 10Gy IR, giving rise to exosomes that were designated as non-damage-derived (NDD EVs) or damage-derived (DD EVs), respectively. Recipient HEK293T cells were incubated with these exosomes for 24h then treated with 10Gy IR or left untreated, and the expression of DNA damage markers, γH2AX and pCHK1, was analysed. Incubation with DD EVs resulted in significantly lower γH2AX and pCHK1 levels in irradiated recipient cells than in those receiving NDD EVs (Figure 1B-D, Figure S2). This reduction in DNA damage markers was also sustained at later time points (Figure S3A and B) suggesting DD EVs lead to radio-resistance via enhanced DNA repair. It has previously been suggested that RNA species can contribute to bystander effects, including enhancement of radio-resistance (Shaw and Gullerova, 2021). Additionally, a similar radioprotective phenotype was shown to be abrogated by RNase A treatment of exosomes (Mutschelknaus *et al*., 2016), suggesting RNA involvement. We therefore hypothesised that YBX1 functions within this process via its ability to specifically load sncRNA species into exosomes. To test this hypothesis, we utilised a YBX1 knockout cell line (YBX1-/-) (Figure S3C and D) and showed that the radioprotective phenotype observed in Figure 1D was diminished when recipient cells were treated with DD EVs from YBX1-/-donor cells (Figure 1E). These data suggest that DNA damage-derived exosomes confer YBX1-dependent radioprotection to bystander cells via enhanced DNA repair.

**Figure 1.**
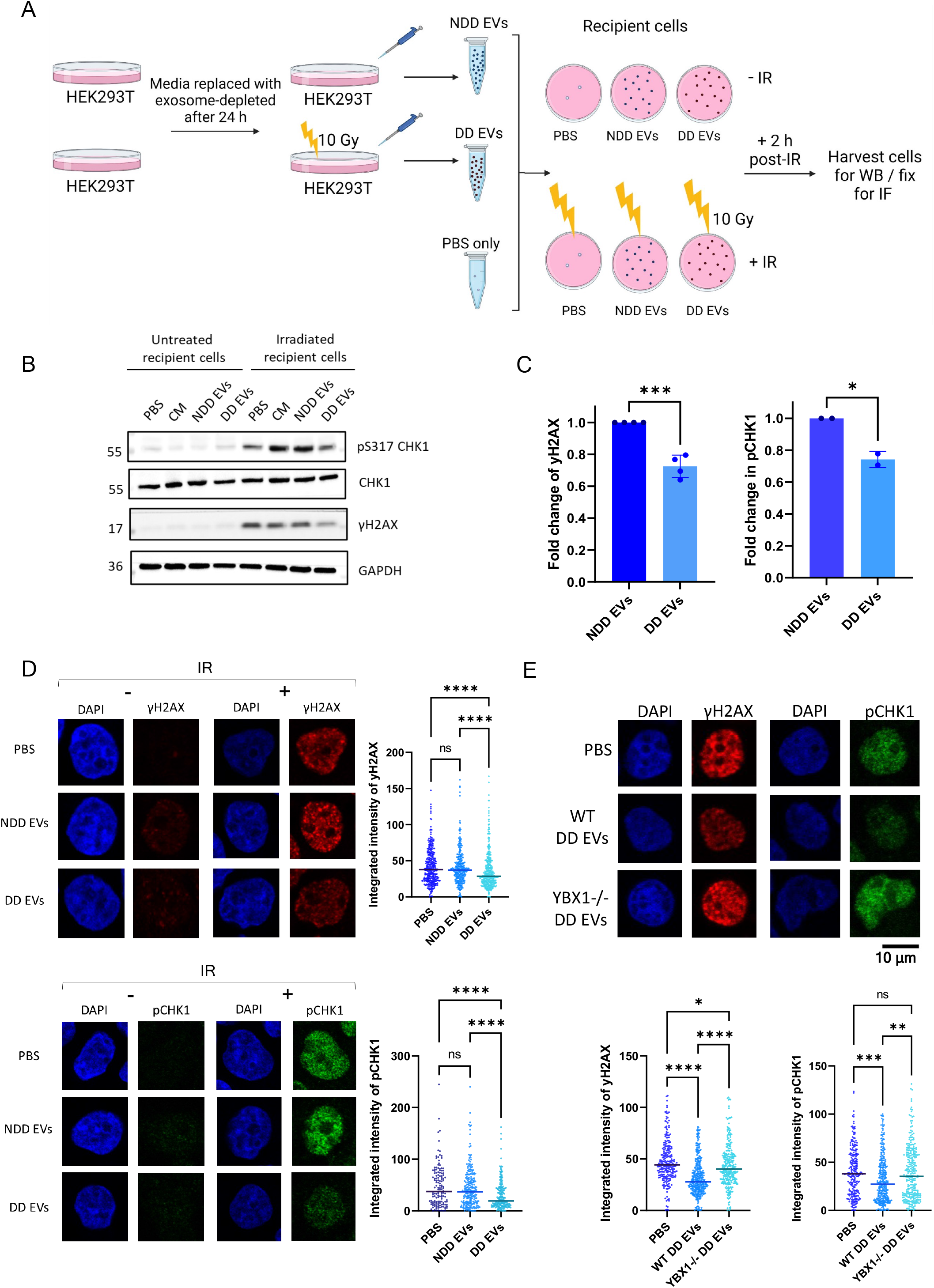
Exosomes derived from damaged cells mediate protective YBX1 dependent bystander effect. A) Schematic overview of exosome transfer experiment, generated in Biorender. NDD EVs = non-damage-derived exosomes, DD EVs = damage-derived exosomes, IR = ionising radiation. B) Representative Western blot showing expression levels of pCHK1 S317 and γH2AX, with CHK1 and GAPDH as loading controls, following exosome transfer and irradiation (10Gy, 2h). C) Bar chart showing the average fold change in γH2AX and pCHK1 expression levels in IR-treated cells upon exosome transfer. Error bars, mean ± SEM. N ≥ 2, independent experiments. Statistical significance was determined using Student’s t-test, *p ≤ 0.05, ***p ≤ 0.001. D) Left panel shows representative immunofluorescence images of γH2AX and pCHK1 expression levels upon transfer of NDD EVs, DD EVs or a PBS control in IR-treated (10Gy, 2h) and untreated cells. Right panel shows quantification from independent experiments where N ≥ 100. Statistical significance was determined using Kruskall-Wallis test with Dunn’s multiple comparison., ****p ≤ 0.0001. E) Top panel shows representative immunofluorescence images of γH2AX and pCHK1 expression levels upon transfer of DD EVs from wild type (WT) or YBX1 knockout (YBX1-/-) donor cells in IR-treated (10Gy, 2h) recipient cells. Bottom panel shows quantification from independent experiments where N ≥ 100. Statistical significance was determined using Kruskall-Wallis test with Dunn’s multiple comparison., ****p ≤ 0.0001.

### YRNA derived fragments in exosomes are responsive to DNA damage and YBX1 KO

Next, we aimed to identify and characterise candidate RNA species which were responsive to both damage and YBX1 knockout conditions and may contribute to the observed phenotype. To do this, we isolated exosomal RNA from wild type (WT) NDD EVs, WT DD EVs and YBX1-/-DD EVs and preformed small RNA sequencing (Figure 2A). Among the identified species of interest were various tRNA-derived fragments and miRNA (Figure S4 and 5), as well as Y RNA-derived fragments (Figure 2B and C). Interestingly, Gene Set Enrichment analysis of the common differentially expressed miRNA species (those that were upregulated upon DNA damage in a YBX1-dependent manner) identified pathways including UV response and G2/M checkpoint (Figure S5E). Similar analysis of the common differentially expressed tsRNA species using the tsRFun database included cell cycle and apoptosis pathways being affected (Figure S4E), both pointing towards possible relevance to radioprotection.

**Figure 2.**
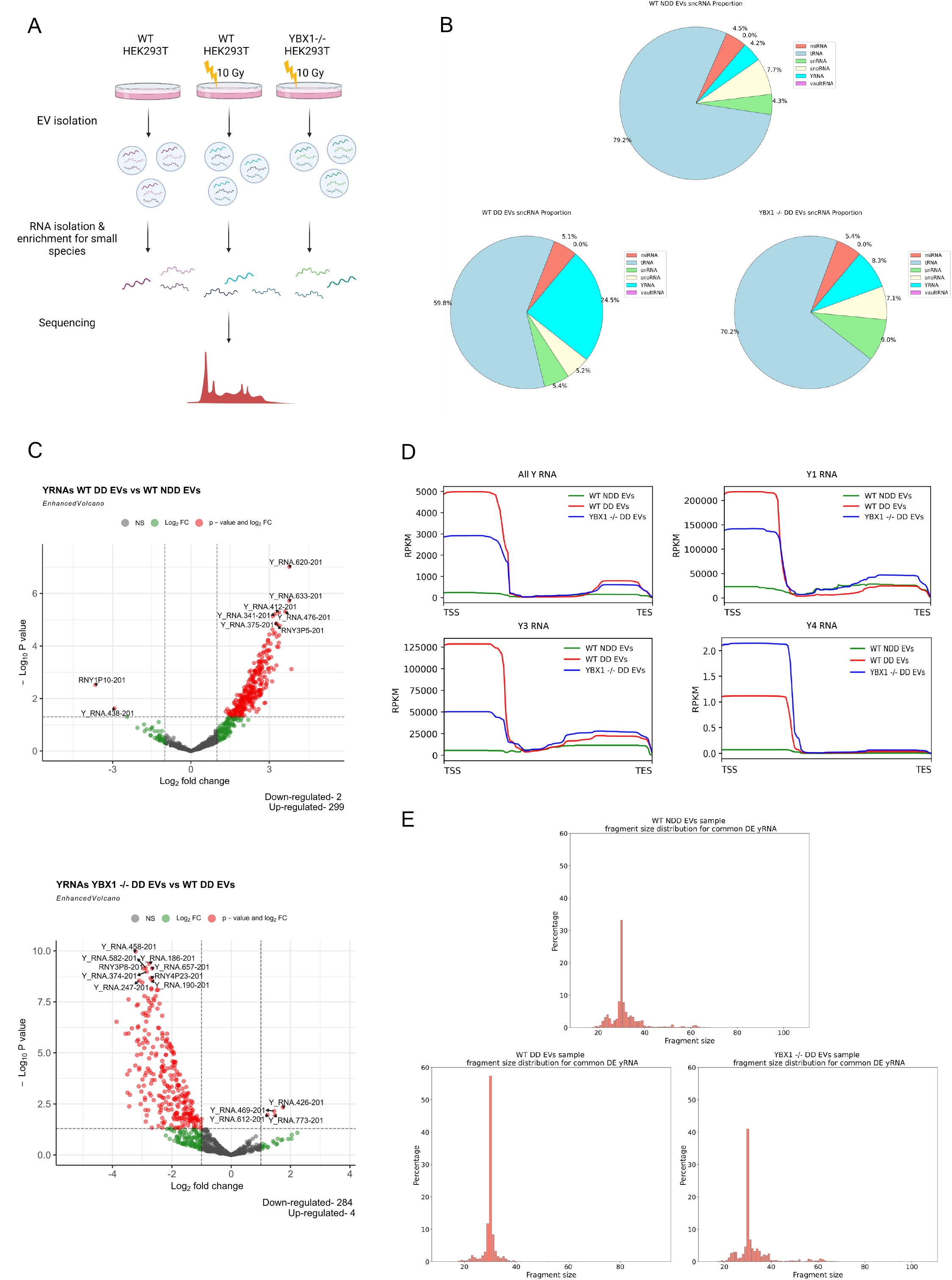
YRNA derived fragments in exosomes are responsive to DNA damage and YBX1 knockout. A) Schematic overview of method followed to obtain RNA from exosomes for small RNA sequencing, generated in Biorender. B) Pie charts showing the proportion of reads corresponding to different classes of small non-coding RNA (sncRNA) obtained from wild-type non-damage-derived exosomes (WT NDD EVs), wild-type damage-derived exosomes (WT DD EVs) and YBX1 knockout damage-derived exosomes (YBX1-/-DD EVs). C) Volcano plots showing differentially expressed Y RNA in exosomes upon damage (top) and YBX1 knockout (bottom). The red points show up-and down-regulated Y RNA-corresponding reads with log2FC > 1 and log2FC < −1 and P-adj < 0.001, respectively. D) Metagene distribution of commonly differentially expressed, Y1, Y3 and Y4 Y RNA reads mapped along normalised gene length, in relation to transcription start and end sites (TSS and TES, respectively). E) Size distribution of reads corresponding to Y RNA from WT NDD EVs, WT DD EVs and YBX1-/-DD EVs from left to right, respectively.

While a large proportion of the reads in our dataset correspond to tRNA fragments (Figure 2B), reads corresponding to Y RNA-derived fragments appeared more highly and specifically responsive to both damage and YBX1 knockout conditions (Figure 2B and C). We therefore decided to focus on Y RNA fragments for further investigation, due to their strong upregulation upon damage in a YBX1-dependent manner. Metagene analysis showed that Y RNA fragments whose presence in exosomes was upregulated upon DNA damage and downregulated upon YBX1 knockout mapped primarily to a region near the transcription start site (TSS) of Y1 and Y3 genes, suggesting these fragments are largely derived from the 5’ end of Y1 and Y3 RNA (Figure 2D). The most enriched fragment length was 30 nt, corresponding to the previously reported ysRNA size of 24-32 nt (Billmeier *et al*., 2022) (Figure 2E and Figure S6). We also show via Northern blot that the levels of Y1 and Y3 RNA are not changing in cells mimicking exosome donor conditions (Figure S7), supporting YBX1 involvement in a packaging capacity, as opposed to influencing Y RNA biogenesis. Overall, these results identified 30 nt 5’ fragments of Y1 and Y3 RNA as candidates for driving the observed radioprotective phenotype.

### YsRNA are nuclear, methylated by NSUN2 and interact with YBX1 at DSBs

The 30 nt sequences derived directly from the 5’ end of Y1 and Y3 genes accounted for 40% and 34% of total Y RNA reads, respectively. We therefore generated synthetic oligonucleotides corresponding to these sequences for further investigation, termed Y1 5’ and Y3 5’, alongside a control 30 nt sequence derived from the 3’ terminus of the Y1 gene (which was not significantly changing in our dataset), termed Y1 3’. In addition to unmodified oligonucleotides, we also generated fluorescently labelled and biotinylated versions of these oligonucleotides, all of which are represented in Figure 3A. First, we sought to determine the subcellular localisation of these ysRNA upon IR treatment in order to assess the likelihood of them influencing DDR. By tracking fluorescently labelled Y1 3’, Y1 5’ and Y3 5’ oligonucleotides via microscopy, we observed significantly higher nuclear localisation of Y3 5’ and, to a lesser extent, Y1 5’ upon IR treatment compared with the control Y1 3’ (Figure 3B and Figure S8A).

**Figure 3.**
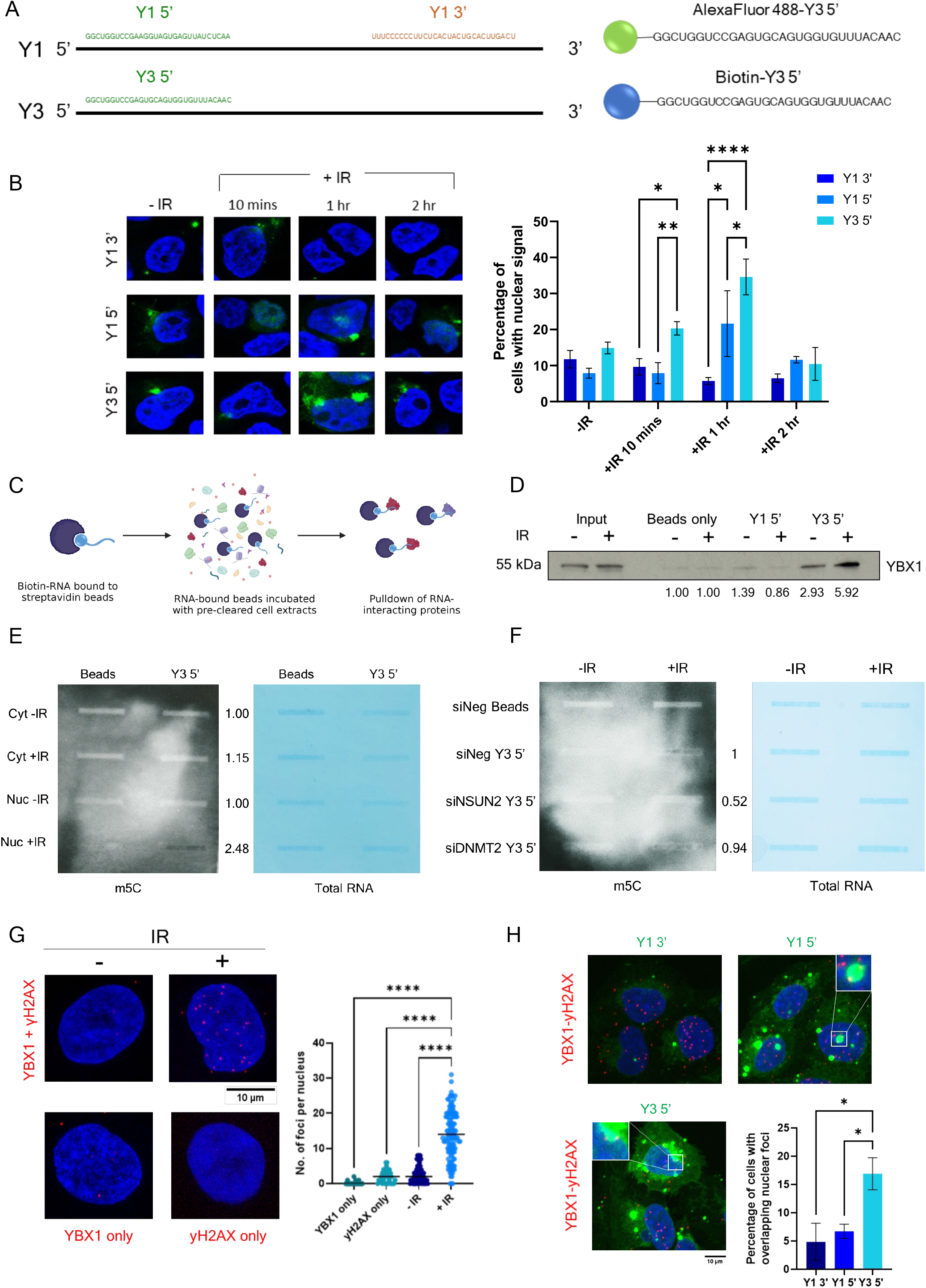
YsRNA are nuclear, modified by NSUN2 and interact with YBX1 at DSBs. A) Representation of synthetic ysRNA oligonucleotide sequences used for subsequent experiments. B) Representative images (left) and quantification (right) of nuclear localisation of AlexaFluor488-labelled ysRNA oligonucleotides following transfection into cells and ionising radiation (IR)-treatment (10Gy, indicated times). Quantification was carried out using ImageJ and represented as the percentage of cells per frame with nuclear fluorescent signal (N ≥ 10 frames). Statistical significance was determined using two-way ANOVA, *p ≤ 0.05, **p ≤ 0.01, ****p ≤ 0.0001. C) Schematic overview of biotinylated RNA immunoprecipitation, generated in Biorender. D) Co-immunoprecipitation of YBX1 with biotinylated Y3 5’ or Y1 5’ ysRNA oligonucleotide, or beads only control (no RNA). Numbers represent fold change in band intensity compared with beads only control, normalised to input. Quantified using ImageJ. E) Slot blot for m5C modification (left) and total RNA stain (right) following pulldown of biotinylated Y3 5’ ysRNA oligonucleotide, or beads only control, after incubation with cytoplasmic (Cyt) and nuclear (Nuc) fractions from IR-treated (10Gy, 10 mins, +IR) and untreated (-IR) cells. Numbers represent signal intensity of m5C in Y3 5’ pulldown, normalised to corresponding total RNA signal. Quantified using ImageJ. F) Slot blot for m5C modification (left) and total RNA stain (right) following pulldown of biotinylated Y3 5’ ysRNA oligonucleotide, or beads only control, after incubation with nuclear fractions from IR-treated (10Gy, 10 mins, +IR) and untreated (-IR) cells, which had been subjected to siRNA-mediated knockdown of NSUN2 or DNMT2 (60nM, 48h) or treated with a negative control siRNA (siNeg). Numbers represent signal intensity of m5C in Y3 5’ pulldown, normalised to corresponding total RNA signal. Quantified using ImageJ. G) PLA of YBX1 and γH2AX, with and without IR treatment (10Gy, +IR and-IR, respectively), including single antibody control experiments. Left panel shows representative images, right panel shows quantification carried out in CellProfiler. N ≥ 100. Statistical significance was determined using Kruskall-Wallis test with Dunn’s multiple comparison. ****p ≤ 0.0001. H) Representative images (top and bottom left) and quantification (bottom right) of PLA between YBX1 and γH2AX, combined with transfection of AlexaFluor488-labelled synthetic ysRNA oligonucleotides (Y1 3’, Y1 5’ and Y3 5’). Quantification represented as percentage of cells with overlapping green and red foci per frame, N ≥ 6 frames. Statistical significance determined by Kruskall-Wallis test with Dunn’s multiple comparison. *p ≤ 0.05.

Next, we aimed to identify associated protein effectors which may help stabilise ysRNA in the nucleus by forming RNPs. YBX1 had previously been identified as an interactor of full length Y3 RNA(Kohn et al., 2015), so we investigated whether YBX1 could also bind 5’ ysRNA in the nucleus of recipient cells. Using synthetic biotinylated ysRNA oligonucleotides, we performed RNA co-immunoprecipitation (Figure 3C) and indeed observe an interaction between Y3 5’ ysRNA and YBX1 which is enhanced upon DNA damage induction (Figure 3D and Figure S8B). As it has previously been shown that YBX1 is a reader of methylation marks, such as NSUN2-deposited m5C(Chen et al., 2019a), we performed a similar pulldown experiment to determine whether Y3 5’ ysRNA could be m5C-modified, which could be important for its recognition and binding by YBX1. Cytoplasmic and nuclear cell fractions were obtained from either untreated or IR-treated cells (Figure S8C), incubated with biotinylated Y3 5’ ysRNA followed by streptavidin pulldown (Figure 3C). However, to check for modification, RNA was eluted from beads instead of protein and analysed by slot blot. Interestingly, we observe that only nuclear fractions from IR-treated cells were able to facilitate m5C modification of Y3 5’ ysRNA, but not cytoplasmic or non-treated nuclear fractions (Figure 3E). We then performed the same experiment using nuclear fractions from IR-treated cells which had been depleted of NSUN2 or DNMT2 by siRNA (Figure S8D) and show that this methylation is dependent on NSUN2 (Figure 3F). Therefore, we suggest that NSUN2-dependent methylation of Y3 5’ ysRNA may be facilitating its interaction with YBX1.

To further investigate whether the YBX1-ysRNA complex has a role in DNA repair, we wanted to determine whether the complex could localise to DSBs. First, we employed a proximity ligation assay (PLA) using antibodies against YBX1 and DSB marker, γH2AX, following IR treatment. This quantitative assay allows for detection of *in vivo* protein-protein interactions(Liu et al., 2024). We observed significantly increased numbers of nuclear PLA foci upon IR treatment (Figure 3G, single antibodies were used as negative controls), demonstrating an interaction between YBX1 and γH2AX and suggesting that YBX1 localises to DSBs. We then combined PLA with transfection of fluorescently labelled ysRNA oligonucleotides (Figure 3A) to determine whether the YBX1-ysRNA complex also localised to DSBs. Indeed, we see a significantly higher overlap between Y3 5’ foci and YBX1-γH2AX PLA foci compared with Y1 5’ and Y1 3’ ysRNA (Figure 3H), suggesting Y3 5’ ysRNA in complex with YBX1 is localised at DSBs. Overall, we show that Y3 5’ ysRNA translocate to the nucleus upon IR, where it becomes methylated by NSUN2 and interacts with YBX1 at DSB sites.

### YsRNA/YBX1 promote efficient DDR and cell survival

In order to confirm that the YBX1-ysRNA complex was physiologically relevant to DSB repair, we interrogated its influence on DDR and cell survival. Using the synthetic ysRNA oligonucleotides described in Figure 3A, we were able to recapitulate the radioprotective phenotype observed by exosome transfer in Figure 1D. Briefly, cells were transfected with relevant unmodified oligonucleotides, incubated for 24h then treated with IR and levels of pCHK1 and γH2AX measured by immunofluorescence. We observe that treating cells with Y3 5’ ysRNA or a combination of Y3 5’ and Y1 5’ ysRNA was able to significantly reduce pCHK1 and γH2AX (Figure 4A and Figure S9A), suggesting ysRNA does play a role in enhancing DNA repair. Conversely, when we employ antisense oligonucleotides (ASOs) against Y3 RNA to downregulate its expression (Figure 4B and Figure S9B), we observe increased levels of γH2AX after IR treatment compared with a luciferase-targeting control ASO (Figure 4C). Together, these data support Y3 5’ ysRNA involvement in DDR following IR exposure. Interestingly, we observe the same trend in γH2AX expression when damage is induced in YBX1-/-cells (Figure 4D), suggesting complementarity in the functions of these two factors.

**Figure 4.**
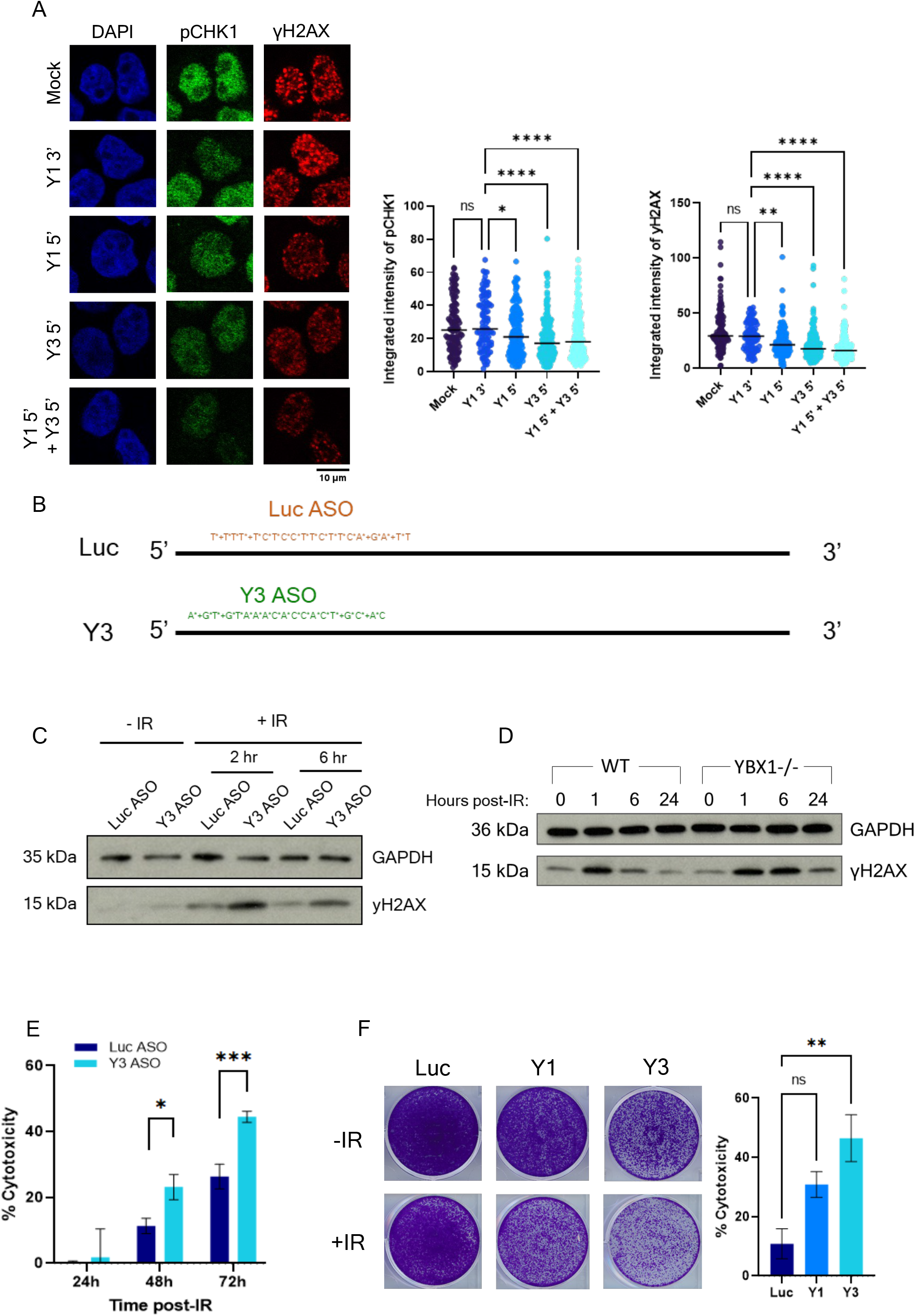
YsRNA/YBX1 promote efficient DDR and cell survival. A) Representative immunofluorescence images (left) and quantification (right) showing levels of pCHK1 S317 and γH2AX following transfection with synthetic ysRNA oligonucleotides, or mock control transfection, and IR-treatment (10Gy). Nuclear signal intensity was quantified using CellProfiler. N ≥ 100. Statistical significance determined by Kruskall-Wallis test with Dunn’s multiple comparison. *p ≤ 0.05, **p ≤ 0.01, ****p ≤ 0.0001. B) Representation of ASO sequences used to downregulate expression of Y3 RNA (Y3 ASO) and luciferase control (Luc ASO). C) Western blot showing expression of γH2AX and GAPDH as a loading control following IR treatment (10Gy, +IR, indicated time points) of cells treated for 48h with control (Luc ASO) or Y3-RNA targeting ASO (Y3 ASO). D) Western blot showing expression of γH2AX, and GAPDH as a loading control, in untreated (0) or IR-treated (10Gy) wild type (WT) or YBX1 knockout (YBX1-/-) cells at various time points after damage (1, 6 and 24h). E) Quantification of MTT assay, showing the cytotoxicity induced by IR (5Gy) in cells treated with Y3 or control (Luc) ASO. Mean ± SEM, N ≥ 12. Statistical significance determined using multiple unpaired t-tests. *p ≤ 0.05, ***p ≤ 0.001. F) Representative images (left) and quantification (right) of clonogenic survival assay. Cells treated with control (Luc), Y1 or Y3 ASO (48h) were untreated (-IR) or subjected to IR (2Gy, +IR) and growth measured after 5 days. Quantification represented as cytotoxicity upon IR treatment. Mean ± SEM. Statistical significance determined by Kruskall-Wallis test with Dunn’s multiple comparison, **p ≤ 0.01.

To ascertain whether this influence on DDR impacted overall cell survival, we performed MTT and clonogenic survival assays in HeLa cells treated with Y3-targeting ASOs (alongside luciferase-targeting and Y1-targeting control ASOs) upon IR treatment. Although there was small reduction in cell survival observed upon Y3 perturbation in untreated cells, relative survival upon IR treatment of Y3 ASO-treated cells was significantly reduced compared with controls (Figure 4E and F). Together, these data suggest that ysRNA, in complex with YBX1, promotes efficient DNA repair and increased cell survival following IR-induced damage.

### YBX1 interacts with PARP1, facilitating ADP-ribosylation of ysRNA

Our next goal was to determine how the YBX1-ysRNA complex influences DDR. As little is known about the function of ysRNA, we decided to focus on known roles of YBX1 in DDR. As previously mentioned, YBX1 has been reported to interact with and promote the activity of PARP1 at DSBs(Alemasova *et al*., 2018; Naumenko *et al*., 2020). We therefore began by confirming this interaction within cells following IR treatment via PLA and co-immunoprecipitation. Indeed, we show increased PLA foci numbers corresponding to YBX1-PARP1 interaction following IR treatment (Figure 5A), which was confirmed by co-immunoprecipitation of overexpressed YFP-tagged PARP1 with YBX1 (Figure 5B). Subsequently, we again combined PLA with fluorescently labelled ysRNA transfection to determine whether ysRNA-YBX1 complexes may also co-localise with PARP1. As hypothesised, we see significantly higher overlap between fluorescently-labelled Y3 5’ ysRNA and YBX1-PARP1 PLA foci in the nucleus than control Y1 3’ ysRNA, suggesting formation of a nuclear ysRNA-YBX1-PARP1 complex (Figure 5C).

**Figure 5.**
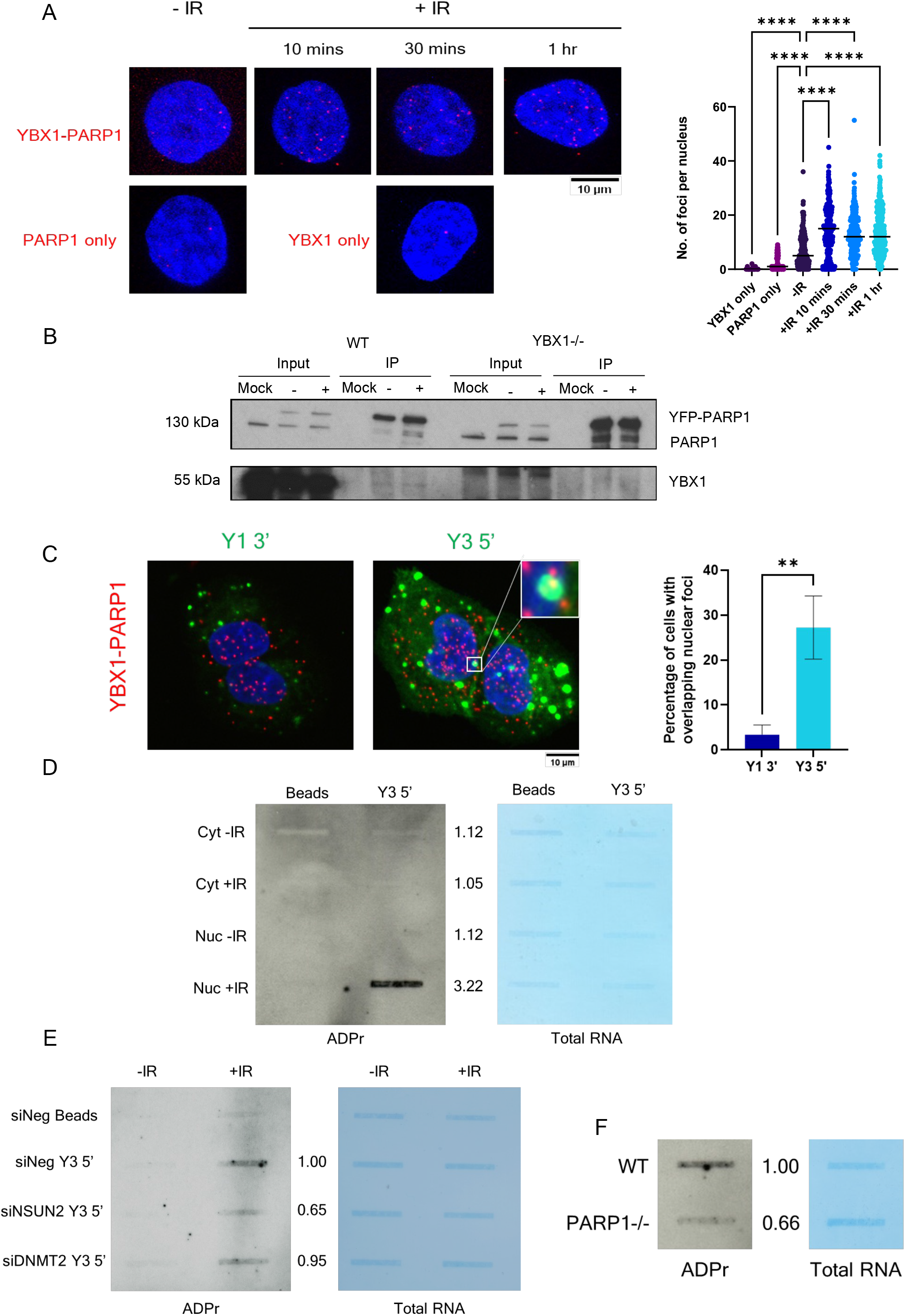
YBX1 interacts with PARP1, facilitating ADP-ribosylation of ysRNA. A) PLA of YBX1 and γH2AX, with and without IR treatment (10Gy, +IR and-IR, respectively), including single antibody control assays. Left panel shows representative images, right panel shows quantification carried out in CellProfiler. Statistical significance was determined using Kruskall-Wallis test with Dunn’s multiple comparison, ****p ≤ 0.0001. B) Detection of YBX1 in co-immunoprecipitation of YFP-PARP1 from mock-transfected or YFP-PARP1-transfected wild type (WT) and YBX1 knockout (YBX1-/-) cells, following subjection to 10Gy IR (+) or left untreated (-). C) Representative images (left) and quantification (right) of PLA between YBX1 and PARP1, combined with transfection of AlexaFluor488-labelled synthetic ysRNA oligonucleotides (Y1 3’ and Y3 5’). Quantification represented as percentage of cells with overlapping green and red foci per frame, N ≥ 6 frames. Statistical significance determined by Mann-Whitney test. **p ≤ 0.01. D) Slot blot for ADP-ribosylation (ADPr) modification (left) and total RNA stain (right) following pulldown of biotinylated Y3 5’ ysRNA oligonucleotide or beads only control after incubation with cytoplasmic (Cyt) and nuclear (Nuc) fractions from IR-treated (10Gy, 10 mins, +IR) and untreated (-IR) cells. Numbers represent signal intensity of ADPr in Y3 5’ pulldown, normalised to corresponding total RNA signal. Quantified using ImageJ. E) Slot blot for ADPr modification (left) and total RNA stain (right) following pulldown of biotinylated Y3 5’ ysRNA oligonucleotide, or beads only control, after incubation with nuclear fractions from IR-treated (10Gy, 10 mins, +IR) and untreated (-IR) cells, which had been subjected to siRNA-mediated knockdown of NSUN2 or DNMT2 or treated with a negative control siRNA (siNeg) for 48h. Numbers represent signal intensity of ADPr in Y3 5’ pulldown, normalised to corresponding total RNA signal. Quantified using ImageJ. F) Slot blot for ADPr modification (left) and total RNA stain (right) following pulldown of biotinylated Y3 5’ ysRNA oligonucleotide, after incubation with nuclear fractions from IR-treated (10Gy, 10 mins, +IR) wild-type (WT) or PARP1 knockout (PARP1-/-) cells. Numbers represent signal intensity of ADPr in Y3 5’ pulldown, normalised to corresponding total RNA signal. Quantified using ImageJ.

YBX1 has previously been reported to become ADP-ribosylated by PARP1 and been suggested to influence its activity through this modification(Alemasova *et al*., 2018). Furthermore, PARP1 has been shown to bind to RNA substrates through its WGR domain *in vitro*(Huambachano et al., 2011). Therefore, we hypothesised that ysRNA within a complex with YBX1 and PARP1 may also become modified. To test this hypothesis, we performed the same subcellular fractionation and ysRNA pulldown experiment described previously (in relation to Figure 3E), this time probing for ADP-ribosylation via slot blot. In line with previous results, we observe a band corresponding to ADP-ribosylated Y3 5’ ysRNA, only in nuclear fractions of IR-treated cells (Figure 5D). Furthermore, this experiment was repeated using nuclear fractions of NSUN2 or DNMT2 knockdown cells to determine whether methylation is required for ADP-ribosylation, possibly by enabling the YBX1-ysRNA interaction. Indeed, we observed a reduction in ADP-ribosylation signal upon NSUN2 knockdown (Figure 5E), suggesting NSUN2-deposited methylation is important for further modification of ysRNA. Finally, we observed a reduction in ADP-ribosylation signal when Y3 5’ ysRNA pulldown was carried out using nuclear fractions from IR-treated PARP1 knockout cells (Figure 5F), suggesting PARP1 is required for ADP-ribosylation of Y3 5’ ysRNA. Together, these data suggest that Y3 5’ ysRNA bound to YBX1 at DSBs is capable of being modified by PARP1 and this is influenced by NSUN2 activity.

### YBX1/ysRNA promote PARP1 residency at DSBs by affecting its auto-modification

It was previously suggested that trans-ADP ribosylation of YBX1 dominates over PARP1 auto-modification within YBX1-PARP1 complexes *in vitro*(Alemasova *et al*., 2018; Naumenko *et al*., 2020). As PARP1 auto-modification correlates with its dissociation from nucleosome-coupled DNA(Sharma et al., 2019), these findings imply that the proposed stimulation of PARP1 by YBX1 may partially result from a longer retention time of active PARP1 on DNA, due to its reduced auto-modification. We therefore used immunoblotting to investigate levels of PARP1 ADP-ribosylation in cells following IR-induced damage, comparing WT with YBX1-/-cells. Interestingly, we see increased levels of PARP1 auto-ADP-ribosylation when YBX1 is lost (Figure 6A and Figure S10), supporting the hypothesis that YBX1 accepting ADP-ribose chains reduces PARP1 auto-modification. In order to establish whether this alteration in PARP1 auto-modification held relevance to its residency at DSBs, we performed PLA using antibodies against PARP1 and γH2AX as a measure of PARP1 localisation at DSBs. At 10 minutes post-IR treatment, we see a significantly higher number of PLA foci compared with untreated control in WT cells, but this is not seen in YBX1-/-cells (Figure 6B). This suggests the presence of YBX1 allows for PARP1 to be maintained in proximity to γH2AX and, by extension, DSBs for longer. Similarly, co-localisation of PARP1 and γH2AX is also reduced in cells treated with Y3 ASO, as observed by a significantly lower number of PLA foci with Y3 ASO treatment compared with control (Figure 6C). Together, these data suggest that YBX1 and Y3 5’ ysRNA work in combination to promote PARP1 residency at DSBs via reduced auto-modification.

**Figure 6.**
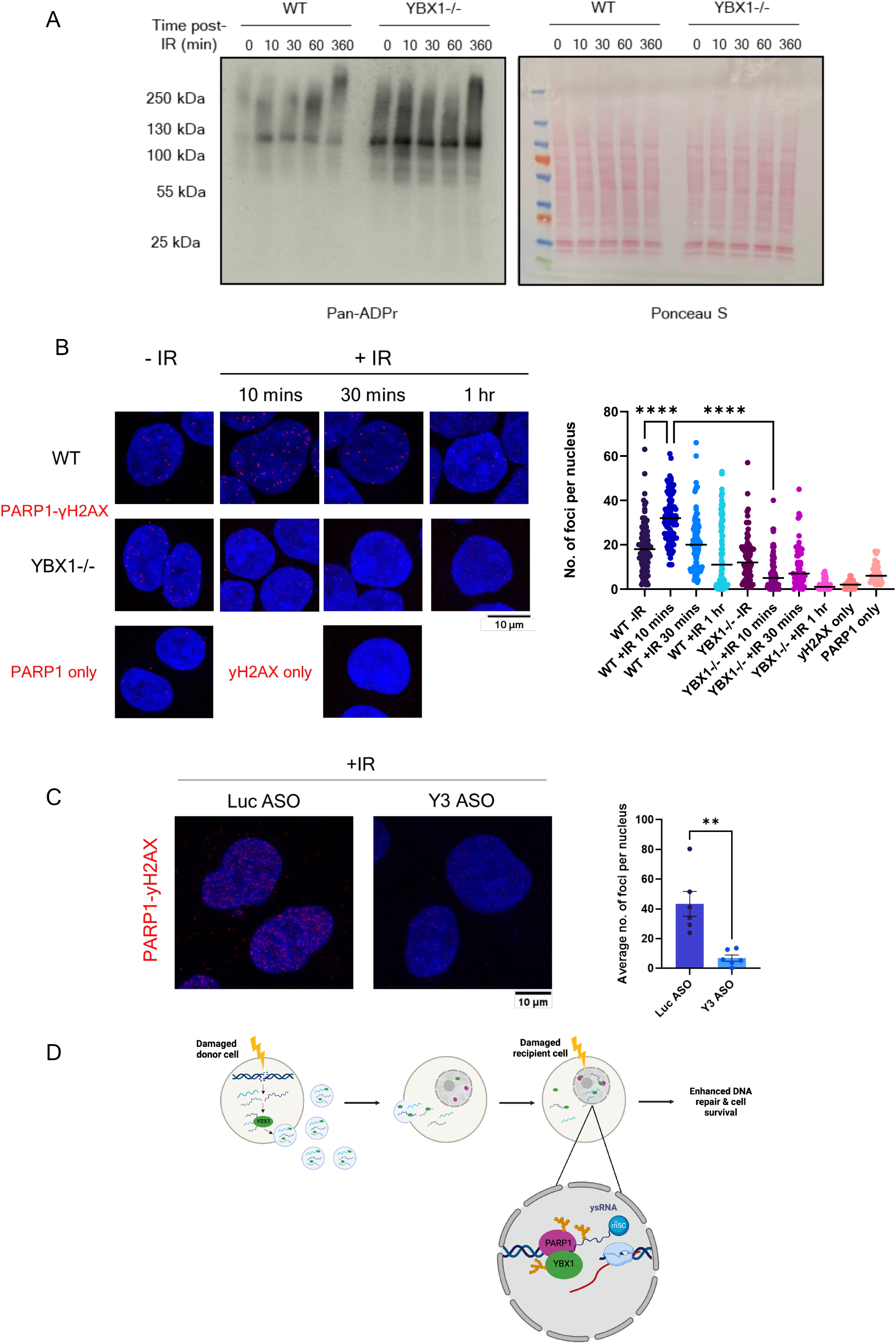
YBX1/ysRNA promote PARP1 residency at DSBs by affecting its auto-modification. A) Western blot for pan-ADP-ribosylation signal (left) in untreated or IR-treated wild-type (WT) or YBX1 knockout (YBX1-/-) cells harvested at indicated time points, alongside total protein levels stained by Ponceau S (right). B) PLA of PARP1 and γH2AX, with and without IR treatment (10Gy, +IR and-IR, respectively), fixed at indicated time points, including single antibody control assays. Left panel shows representative images, right panel shows quantification carried out in CellProfiler. Statistical significance was determined using Kruskall-Wallis test with Dunn’s multiple comparison, ****p ≤ 0.0001. C) PLA of PARP1 and γH2AX upon IR treatment (10Gy, 10 mins) of Y3 or control (Luc) ASO-treated cells (48h). Left panel shows representative images, right panel shows quantification carried out in ImageJ. Mean ± SEM. Statistical significance was determined using Student’s t-test, **p ≤ 0.01. D) Proposed model of YBX1-ysRNA mediated radioprotective bystander phenotype. Generated in Biorender.

## Discussion

Radioprotective bystander effects transmitted by exosomes have been previously studied, largely in terms of their influence on cell survival(Mrowczynski *et al*., 2018; Mutschelknaus *et al*., 2016). While one study did suggest RNA involvement and enhanced DNA repair as a potential mechanism, they did not identify any specific candidate RNA species which may be contributing to this phenotype(Mutschelknaus *et al*., 2016). Here, we show that radioprotection is YBX1-dependent and is driven, at least partially, by Y3-derived ysRNA. It is possible that other species of sncRNA which were identified in our sequencing dataset but not explored (e.g. miRNA or tRNA-derived fragments) may further enhance radioprotection. However, we show that similar reductions in DNA damage markers could be induced by both ysRNA and DD EVs, alongside an influence on overall cell survival, supporting the significant involvement of ysRNA.

Furthermore, we show that Y3-derived ysRNA become localised to the nucleus specifically upon IR-induced DNA damage. One possible mechanism through which this may occur is via binding to Ro60 and the Zipcode-Binding Protein, ZBP1. It has been shown that nuclear localisation of Ro60/Y3 RNA complexes upon UV irradiation is dependent on ZBP1 binding(Sim et al., 2012). Although this work focuses on full length Y3 RNA, as opposed to ysRNA, it may still provide mechanistic insight into ysRNA nuclear localisation and its subsequent role in DDR. Interestingly, both Ro60 and ZBP1 could also be co-purified with YBX1(Sim *et al*., 2012), suggesting proximity between YBX1 and Y3 RNA upon nuclear entry. Nevertheless, this would not explain the relevance of our observed NSUN2-deposited methylation marks on Y3-derived ysRNA, nor support our hypothesised link between methylation and YBX1-ysRNA binding through YBX1 activity as an m5C reader(Chen *et al*., 2019a). As a result, further investigation would be required to delineate the importance of NSUN2-dependent ysRNA methylation in the nucleus.

Our study demonstrates that ysRNA involvement in radioprotection and, by extension, canonical DNA repair, is dependent on YBX1. While it is a known contributor to DNA repair, the exact mechanism by which YBX1 becomes recruited to DNA damage sites remains unknown. It has been suggested by multiple studies to be dependent on RNA molecules(Liu et al., 2016; Nothen et al., 2023), although YBX1 is also known to interact with many other DDR-associated proteins which may influence its chromatin recruitment(Kim *et al*., 2013). Moreover, YBX1 contains an intrinsically disordered domain at its C-terminus and has been reported to undergo liquid-liquid phase separation(Liu *et al*., 2021). Another group suggested YBX1 to participate in phase transitions initiated by poly(ADP-ribose) chains, potentially enabling its DNA repair function(Alemasova and Lavrik, 2017). This is supported by other studies proposing phase separation to occur at DSB sites and contribute to efficient repair(Long et al., 2023; Pessina et al., 2019; Wang et al., 2022). Therefore, it is possible that the association we observe between Y3-derived ysRNA, YBX1 and PARP1 may result from a similar phase separation event, rather than a direct interaction and complex formation.

PARP1 and deposited ADP-ribose chains are known to play important roles in DNA repair(Ray Chaudhuri and Nussenzweig, 2017). We demonstrate, for the first time, PARP1-dependent ADP-ribosylation of ysRNA. While ADP-ribosylation of RNA has been previously reported, this was attributed to other members of the PARP family, such as PARP10, PARP11 and PARP15(Munnur et al., 2019). When tested in cells, RNA ADP-ribosylation was found to be responsive to various stressors, including H_2_O_2_ (a known activator of PARP1 activity), although this did not appear to impact modification of small RNA populations(Weixler et al., 2022). Other proposed roles of RNA ADP-ribosylation include initiation of immune responses and influencing mRNA stability(Groslambert et al., 2021; Munnur *et al*., 2019). PARP1 has been implicated in ADP-ribosylation of phosphorylated DNA ends, which was suggested to potentially promote DNA repair(Groslambert *et al*., 2021), although further investigation is required to determine the exact functions and importance of nucleic acid ADP-ribosylation. Nevertheless, the observed ysRNA ADP-ribosylation within PARP1-YBX1 complexes, and the resulting decrease in PARP1 auto-modification, may result in increased chromatin residency of PARP1. This suggests a previously unknown contribution of ysRNA to DSB repair.

Overall, we propose a model whereby YBX1 plays a dual role in mediating radioprotective bystander responses via Y3-derived ysRNA. First, YBX1 packages ysRNA into exosomes which can be transmitted to bystander cells and enhance DNA repair and cell survival in response to subsequent radiation insult. Secondly, YBX1 binds methylated ysRNA in the nucleus of these cells, facilitating complex formation with PARP1. This allows for ADP-ribosylation of ysRNA and YBX1, reducing PARP1 auto-modification and leading to increased PARP1 residency on chromatin to facilitate efficient DNA repair (Figure 6d). These results provide important insights for a greater understanding of RNA dependent DDR and provide a new perspective on regulation of PARP1 activity, which is highly relevant to cancer therapy.

## Acknowledgements

This work was supported by the Senior Research Fellowship by Cancer Research UK [grant number BVR01170], EPA Trust Fund [BVR01670], and Lee Placito Fund awarded to M.G.

We are grateful to all members of MG lab for their support during the study. We would like to thank Matthew Wood’s Lab for help with exosomes purifications, to Randy Schekman’s Lab for YBX1-/-cells, Ivan Ahel’s Lab for PARP1-/-cells and helpful discussions about PARP1 activity, and to Meliti Skouteri for generation of initial data.

## Author contributions

A.S. performed all experiments and wrote the manuscript. K.A. performed bioinformatic analysis of sRNA-seq. M.G. supervised and designed the project and edited the manuscript. All the authors reviewed and approved the final version of the manuscript.

## Figure legends

**Supplementary Figure 1.**
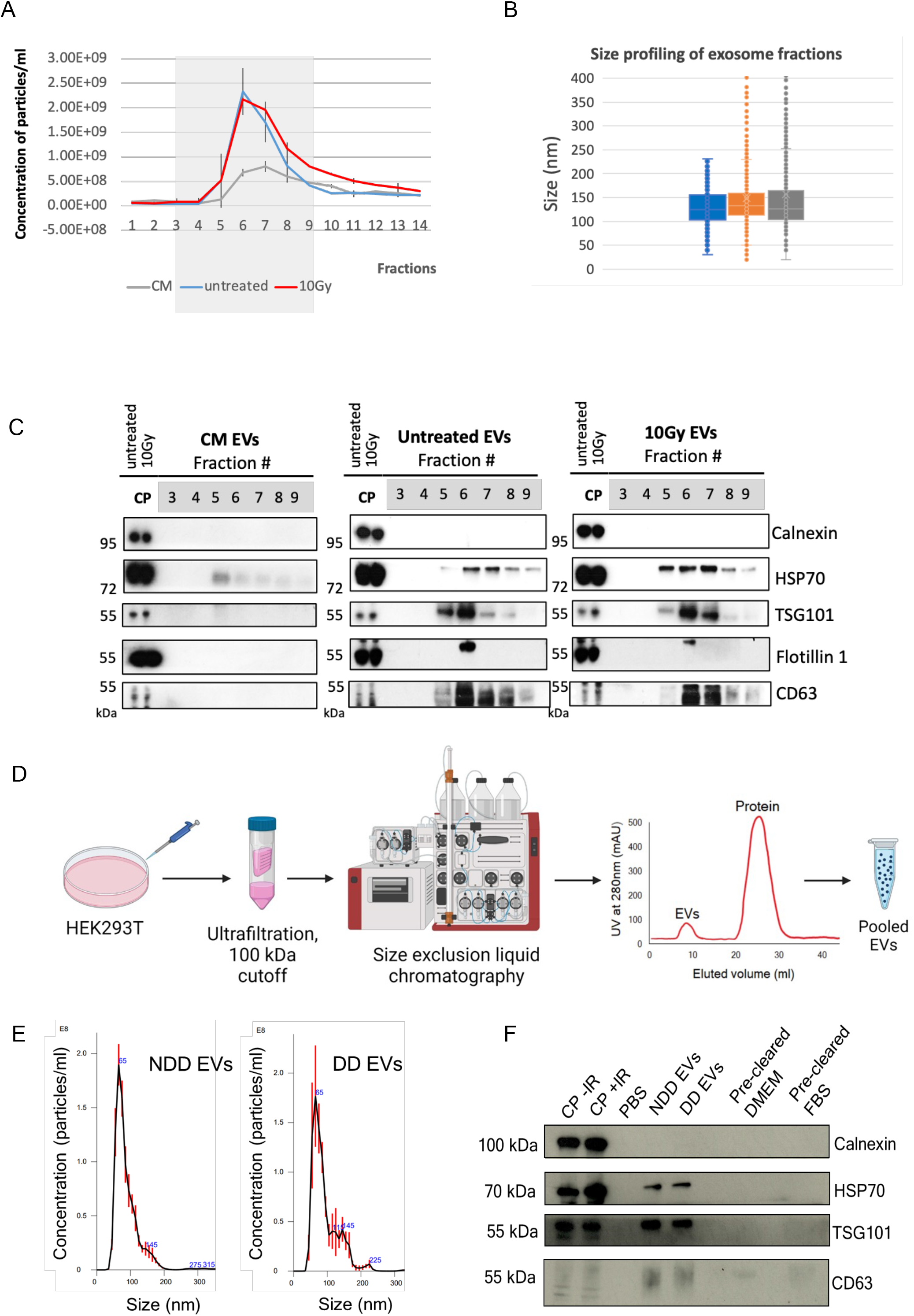
A) Concentration of particles in fractions obtained by size exclusion chromatography, measured by Nanoparticle Tracking Analysis. B) Size distribution of particles in fractions obtained by size exclusion chromatography, measured by Nanoparticle Tracking Analysis. C) Quality control Western blot showing presence of exosome-specific markers, HSP70, TSG101, Flotillin 1 and CD63, and absence of cellular marker, Calnexin, in fractions obtained by size exclusion chromatography and cellular protein (CP) control. D) Workflow for automated size exclusion chromatography, generated in Biorender. E) Concentration and size distribution of particles in pooled EV samples from non-damage-derived (NDD EVs) and damage-derived exosomes (DD EVs) obtained by automated size exclusion chromatography. F) Quality control Western blot showing presence of exosome-specific markers, HSP70, TSG101 and CD63, and absence of cellular marker, Calnexin, in pooled EV samples obtained by size exclusion chromatography, alongside cellular protein (CP), pre-cleared DMEM and pre-cleared FBS controls.

**Supplementary Figure 2.**
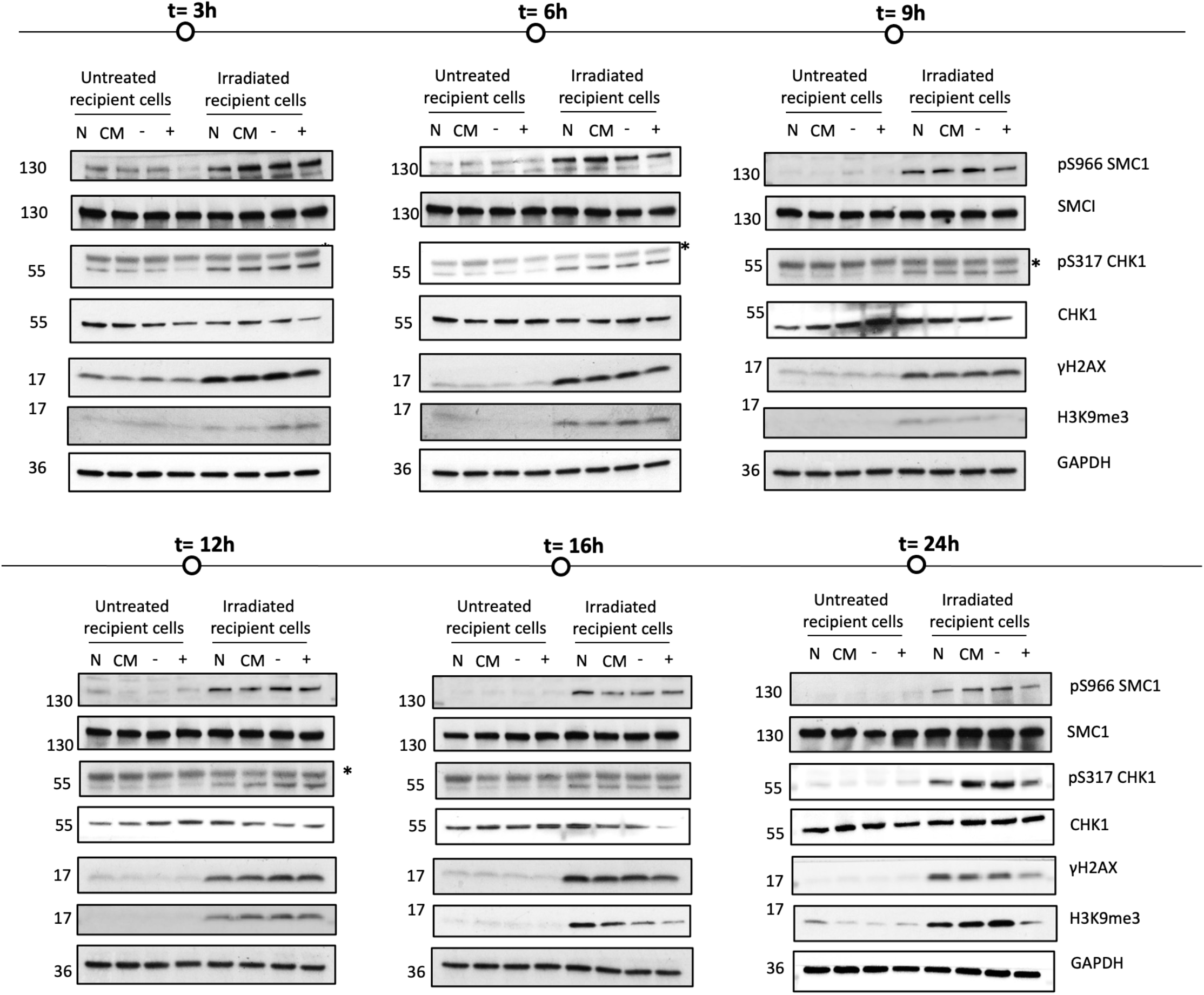
Western blots showing optimisation of time (t) for exosome incubation following transfer. N = negative control, PBS incubation; CM = clear media only;-= non-damage-derived exosomes; + = damage-derived exosomes.

**Supplementary Figure 3.**
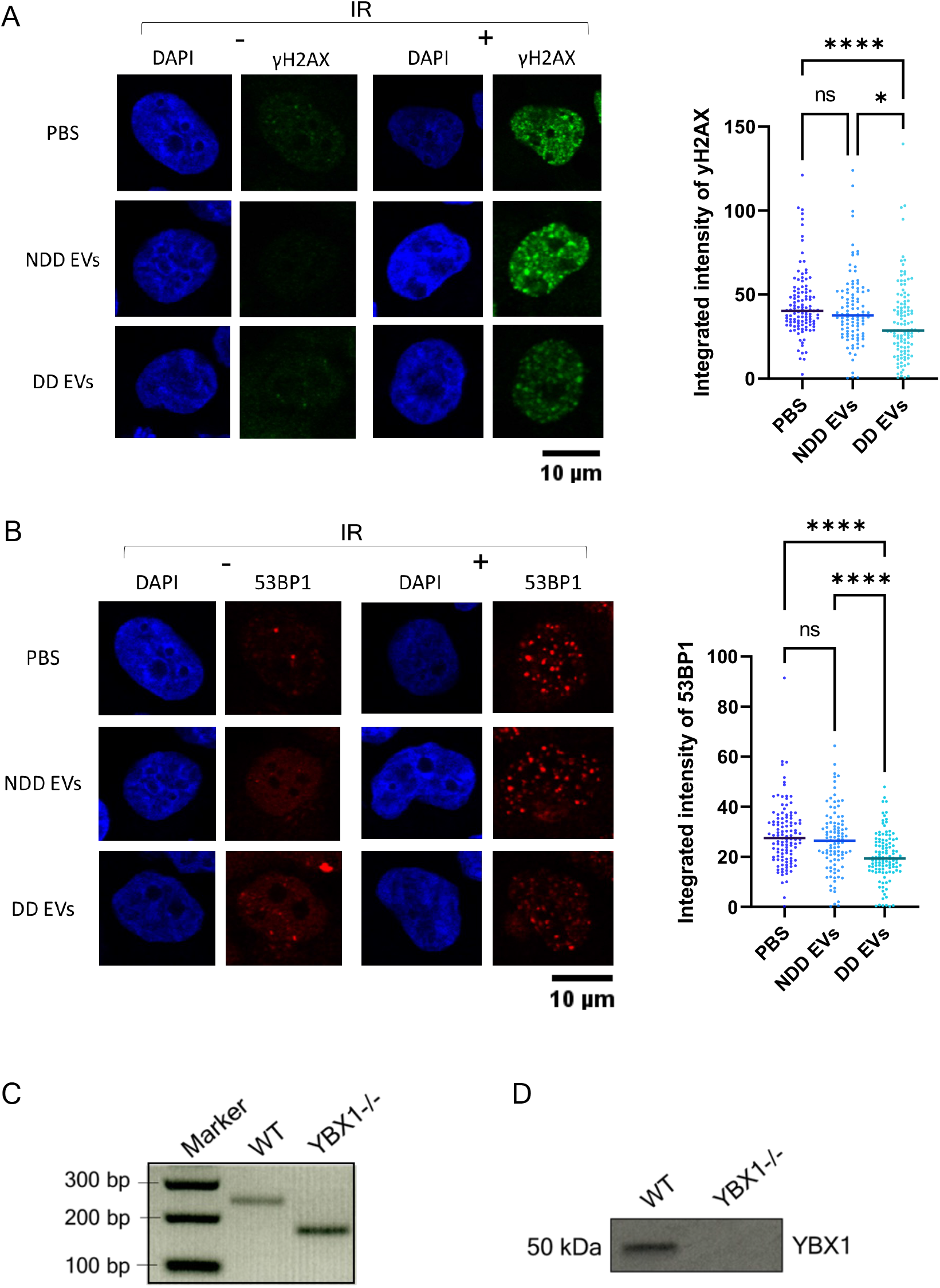
A) Representative immunofluorescence images (left) and quantification (right) showing levels of γH2AX following transfer of non-damage-derived exosomes (NDD EVs), damage-derived exosomes (DD EVs) or a PBS control in IR-treated (10Gy, 6h) and untreated cells. Nuclear signal intensity was quantified using CellProfiler, N ≥ 100. Statistical significance determined by Kruskall-Wallis test with Dunn’s multiple comparison. *p ≤ 0.05, ****p ≤ 0.0001. B) Representative immunofluorescence images (left) and quantification (right) showing levels of 53BP1 foci following transfer of NDD EVs, DD EVs or a PBS control in IR-treated (10Gy, 6h) and untreated cells. Nuclear signal intensity was quantified using CellProfiler, N ≥ 100. Statistical significance determined by Kruskall-Wallis test with Dunn’s multiple comparison, ****p ≤ 0.0001. C) Analysis of wild-type (WT) and YBX1-/-HEK293T cells by PCR flanking the genomic target site of CRISPR-Cas9-mediated editing to confirm knockout. D) Western blot confirming YBX1 knockout.

**Supplementary Figure 4.**
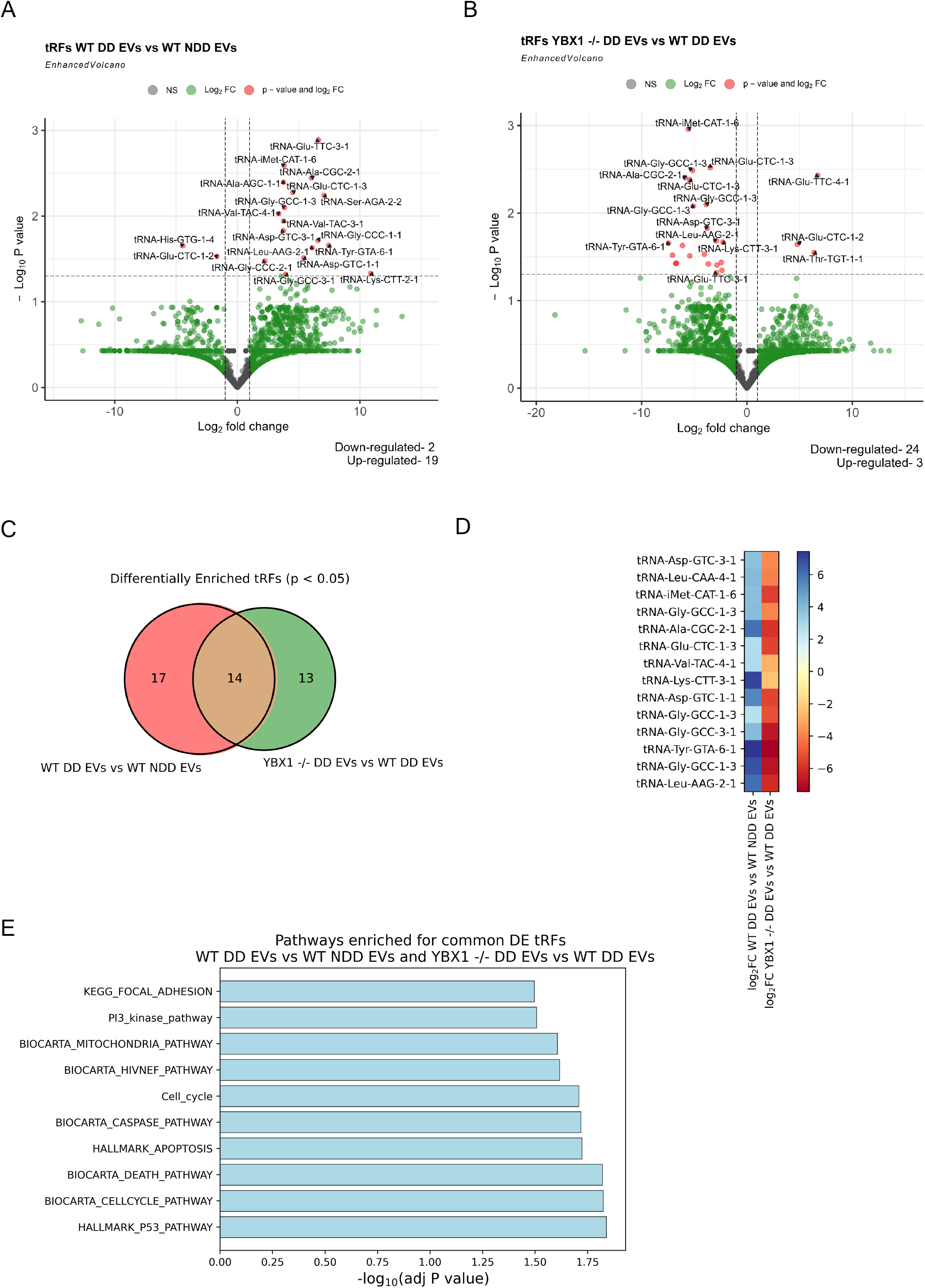
A) Volcano plots showing differentially expressed tRNA-derived fragments (tRFs) in exosomes upon damage (left) and YBX1 knockout (right). The red points show up-and down-regulated Y RNA-corresponding reads with log2FC > 1 and log2FC < −1 and P-adj < 0.001, respectively. B) Venn diagram showing the number of commonly differentially expressed tRFs between damage and YBX1 knockout conditions. C) Heat map of common differentially expressed tRFs identified in b), showing their log2FC in damage vs non-damage-derived exosomes, and YBX1 knockout vs wild-type damage-derived exosomes, respectively. D) Pathways identified by Geneset enrichment analysis as affected by the differentially expressed tRFs whose presence in exosomes is damage and YBX1-dependent.

**Supplementary Figure 5.**
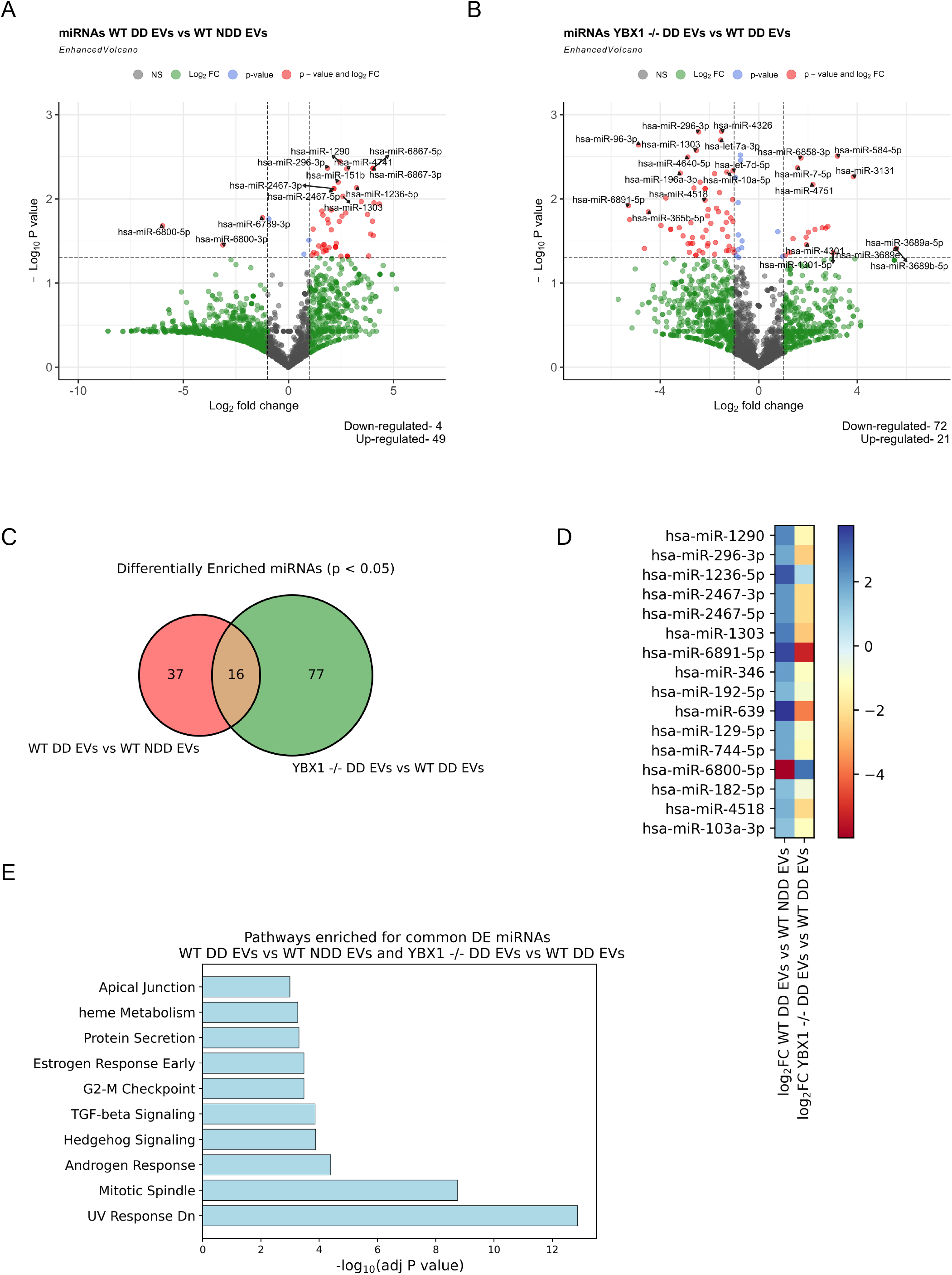
A) Volcano plots showing differentially expressed miRNA species in exosomes upon damage (left) and YBX1 knockout (right). The red points show up-and down-regulated Y RNA-corresponding reads with log2FC > 1 and log2FC < −1 and P-adj < 0.001, respectively. B) Venn diagram showing the number of commonly differentially expressed miRNA species between damage and YBX1 knockout conditions. C) Heat map of common differentially expressed miRNA species identified in b), showing their log2FC in damage vs non-damage-derived exosomes, and YBX1 knockout vs wild-type damage-derived exosomes, respectively. D) Pathways identified by Geneset enrichment analysis as affected by the differentially expressed miRNA species whose presence in exosomes is damage and YBX1-dependent.

**Supplementary Figure 6.**
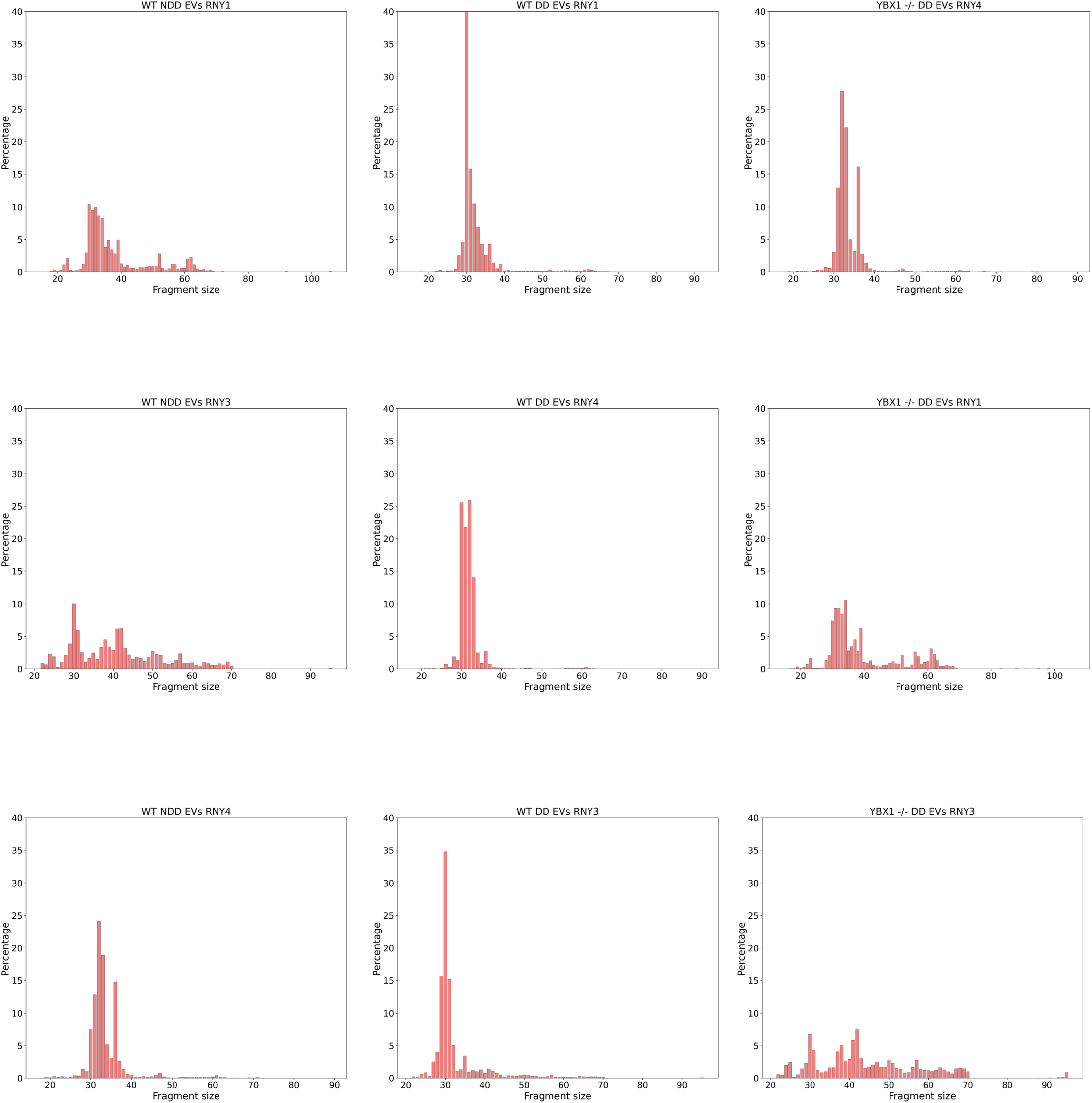
Size distribution of reads corresponding to Y RNA genes (Y1, Y3 and Y4) from WT NDD EVs, WT DD EVs and YBX1-/-DD EVs from left to right, respectively.

**Supplementary Figure 7.**
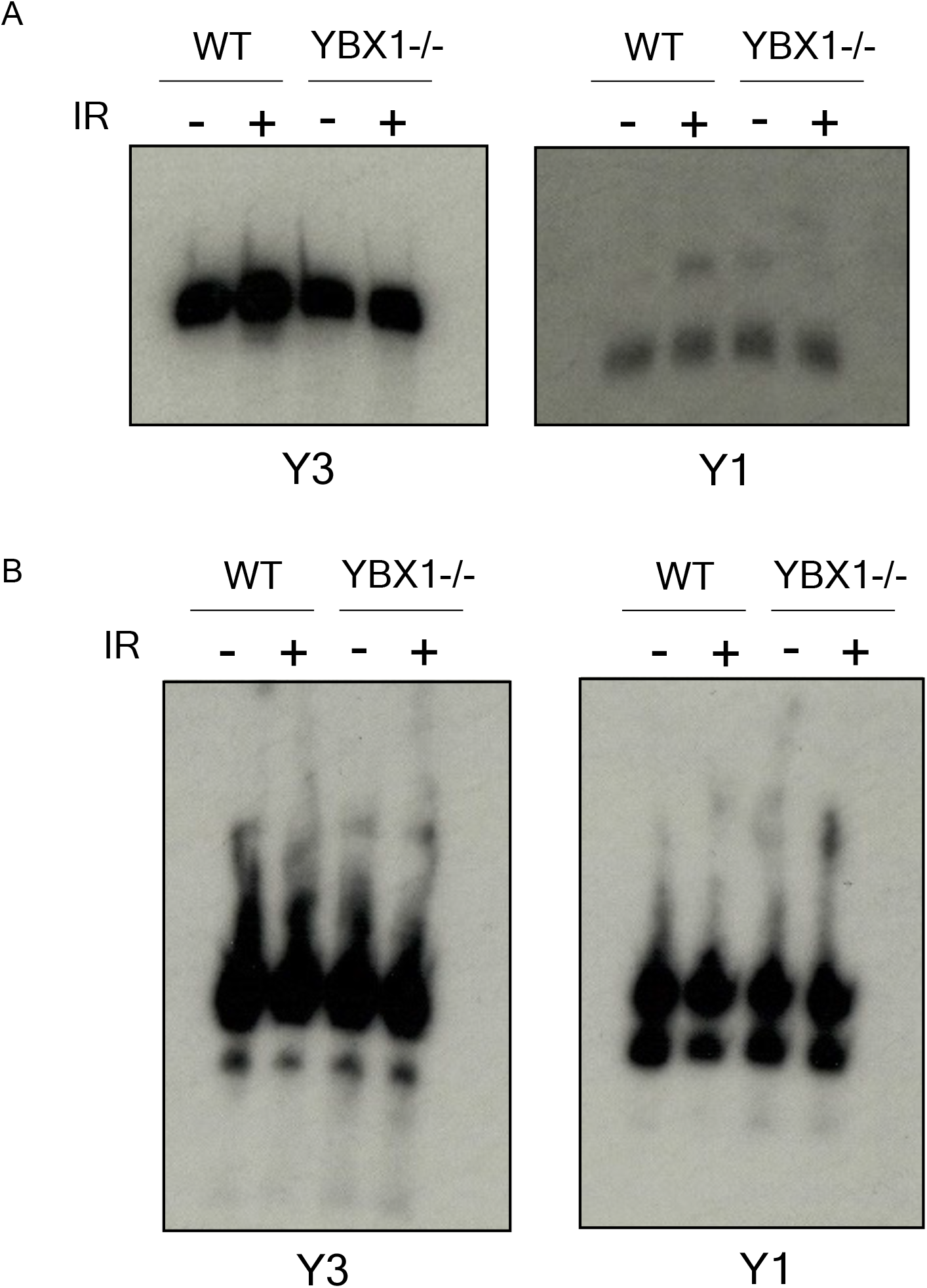
A) Northern blot showing levels of Y3 and Y1 RNA in wild-type (WT) and YBX1 knockout (YBX1-/-) cells which were untreated (-) or subjected to 10Gy IR for 48h (+). B) Northern blot showing levels of Y3 and Y1 RNA and their associated degradation products in wild-type (WT) and YBX1 knockout (YBX1-/-) cells which were untreated (-) or subjected to 10Gy IR for 48h (+).

**Supplementary Figure 8.**
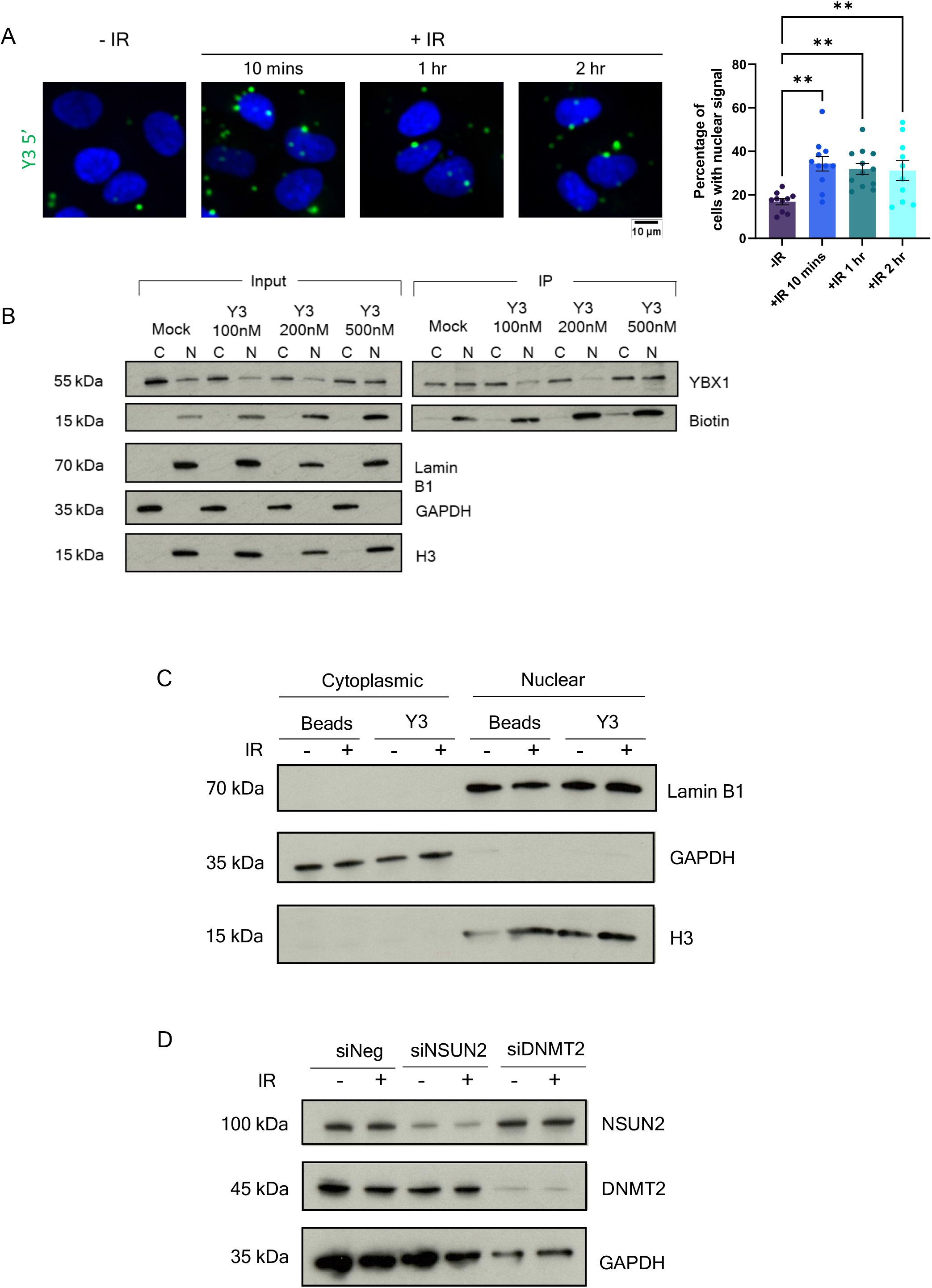
A) Representative images (left) and quantification (right) of nuclear localisation of AlexaFluor488-labelled Y3 5’ ysRNA oligonucleotides following transfection into cells and ionising radiation (IR)-treatment (10Gy, indicated times). Quantification was carried out using ImageJ and represented as the percentage of cells per frame with nuclear fluorescent signal (N ≥ 10 frames). Mean ± SEM. Statistical significance was determined using one-way ANOVA with Dunnett’s multiple comparison, **p ≤ 0.01. B) Western blot showing co-immunoprecipitation (IP) of YBX1 from cytoplasmic (C) and nuclear (N) fractions from cells transfected with varying concentrations of biotinylated Y3 5’ ysRNA synthetic oligonucleotide. Input also includes quality control for fractionation efficiency. C) Western blot for cytoplasmic marker, GAPDH, and nuclear markers, Lamin B1 and histone H3, to check efficiency of fractionation in samples used for RNA immunoprecipitation and slot blot.

**Supplementary Figure 9.**
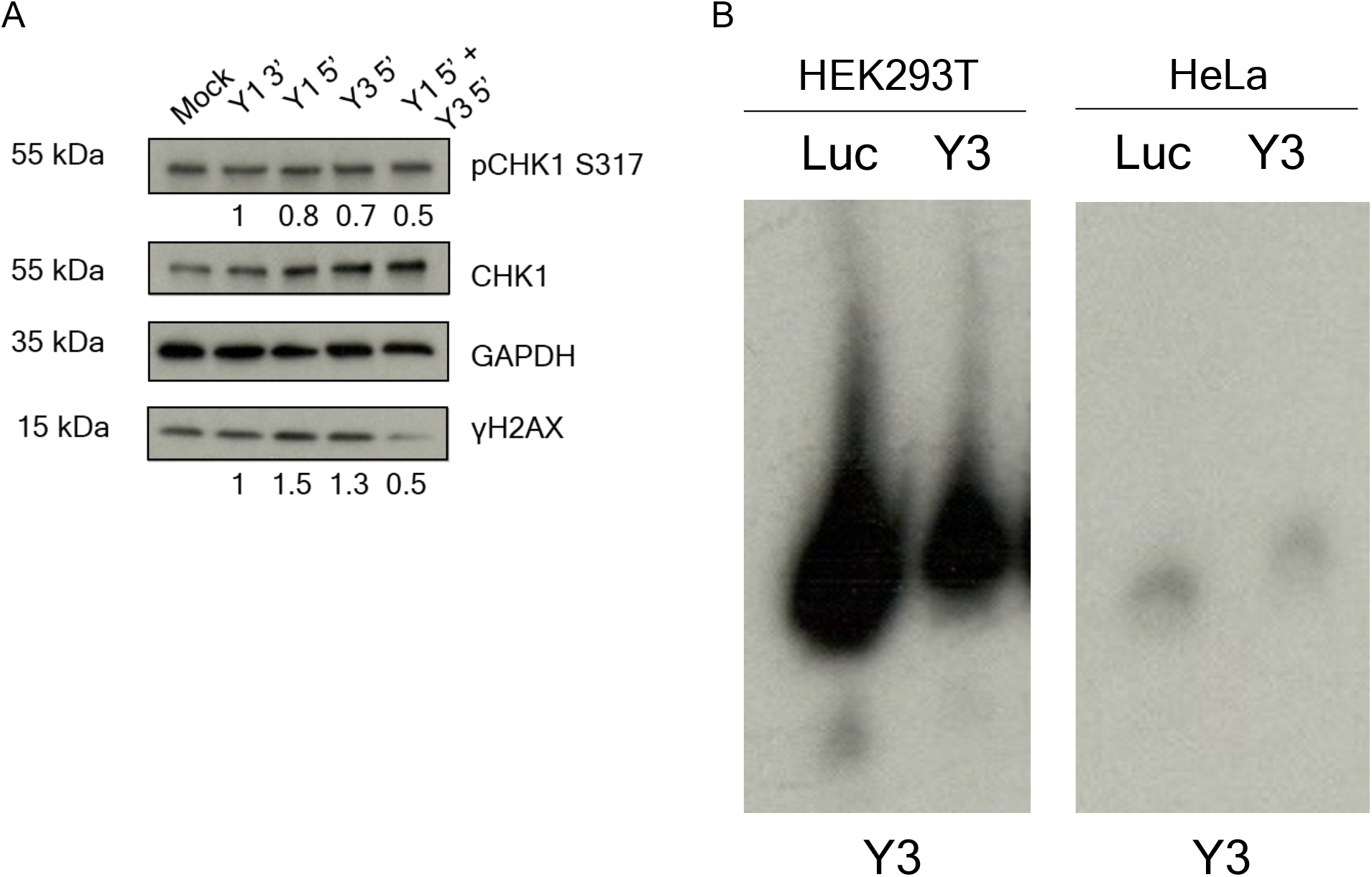
A) Western blot showing expression levels of pCHK1 S317 and γH2AX, with CHK1 and GAPDH as loading controls, following transfection with various synthetic ysRNA oligonucleotides (Y1 3’, Y1 5’, Y3 5’ or a combination of Y1 5’ and Y3 5’, 10nM for 24h) or mock transfected control upon IR treatment (10Gy, 2h). Numbers represent fold change in normalised band intensity over control, quantified using ImageJ. B) Northern blot showing Y3 RNA levels upon control (Luc) and Y3 ASO treatment (125nM, 48h) in HEK293T and HeLa cells.

**Supplementary Figure 10.**
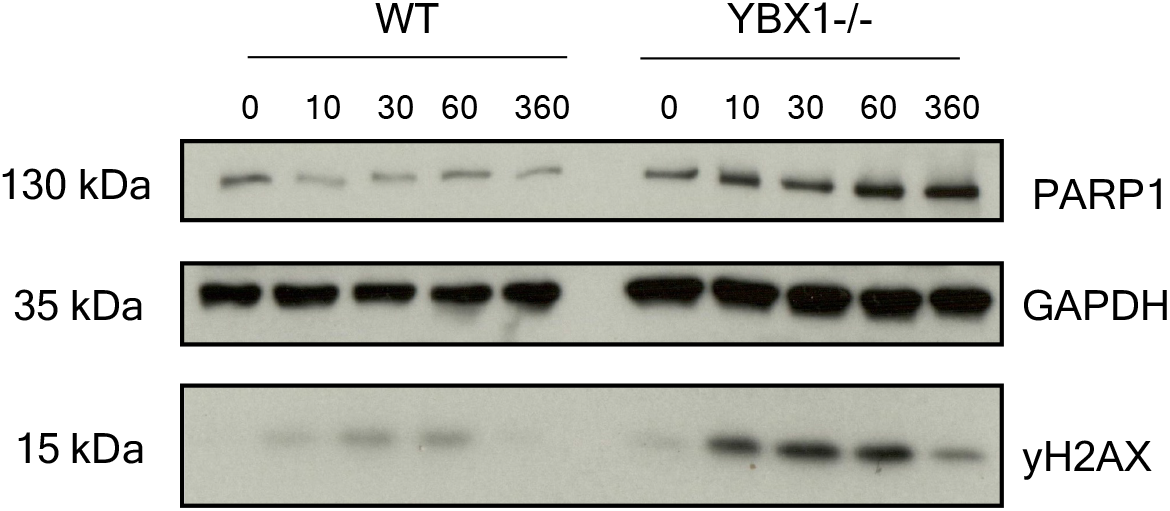
Western blot showing levels of PARP1 and γH2AX, with GAPDH as a loading control, following IR treatment (10Gy) at indicated time points (mins) of wild-type (WT) or YBX1 knockout (YBX1-/-) cells.

## STAR★Methods

### Resource availability

#### Lead contact

Further information and requests for resources and reagents should be directed to and will be fulfilled by the Lead Contact, Monika Gullerova.

### Materials availability

Reagents generated in this study can be made available on request.

### Data and code availability

- Data reported in this paper can be shared by the lead contact upon request.
- Exosome Small RNA-seq data have been deposited to GEO and can be accessed under GSE269346 with private token: kzklgeiitlmptkd
- This paper does not report original code.
- Any additional information required to re-analyse the data reported in this work paper is available from the lead contact upon request.

## Experimental model and study participant details

### Cell lines and cell culture

HeLa, HEK293T, YBX1-/-HEK293T (gifted from the Schekman Lab) and PARP1-/-HEK293T (gifted from the Ahel Lab) cells were maintained at 37°C and 4% CO_2_ in DMEM high glucose medium (Life Technologies #31966047), supplemented with 10% fetal bovine serum (FBS) (Sigma-Aldrich #F9665), 2mM L-glutamine (Life Technologies #25030024) and 100 units/ml penicillin-streptomycin solution (Life Technologies #15140122). TrypLE Express was used to passage cells once 70% confluency reached.

Ionising radiation was delivered at indicated doses (2, 4, 5 or 10 Gy) by source exposure using GRAVITRON RX 30/55.

### FBS pre-clearing

FBS was filtered through a 0.2µm filter then aliquoted into 70ml Polycarbonate Bottles for ultracentrifugation. Centrifugation was carried out using Beckman XPN-80 and type 45 Ti rotor at 4°C and 100,000g for 20h. Clear fraction was collected, filtered and stored at-20°C.

### Exosome isolation

HEK293T cells were seeded into 10x 15 cm dishes per condition. After 24h, cell culture medium was replaced with DMEM + 10% exosome-depleted FBS and cells were exposed to 10 Gy IR, if applicable. After a further 48h, culture medium was extracted and centrifuged at 300g for 5 mins to pellet any cells, centrifuged at 16,500g for 20 mins to pellet microvesicles, and filtered through a 0.22μm PES filter (Corning). Media was then concentrated to 15ml using a 100kDa MWCO tangential flow membrane (Sartorius Vivaflow 50R), followed by further concentration to 2ml by centrifugation at 4,000g for ∼15 mins using 100 kDa concentrators (Fisher #28-9323-63).

2ml samples were loaded into AKTA pure system with UV flow cell (GE Healthcare), connected to a Sepharose 4 fast flow gel-filtration column (GE Healthcare, #170149001) in DPBS, via a 2ml loop. UNICORN software was used to set flow rate to 0.5ml/min with 500μl fraction collection and no protein collection, and to visualise absorbance at 280nm. EV fractions were collected based on UV trace and pooled. Further concentration to 200μl was achieved by centrifugation at 4,000g using 100 kDa centrifugal filters (Merck #UFC210024).

### Nanoparticle Tracking Analysis

Exosome concentration and size distribution measured using NanoSight NS500 (Malvern) and NanoSight 3.2 software. Concentrated, pooled EVs were diluted 1:500 in PBS. 5x measurements were recorded for 10s each.

### Exosome transfer

HEK293T cells were seeded at 20% confluency into 6cm dishes for western blotting, or onto poly-L-lysine coated glass coverslips for immunofluorescence. After 24h, exosome-depleted DMEM, combined with NDD EVs, DD EVs or an equal volume of PBS, replaced the culture media of 2x sets of recipient cells. After a further 24h, one set of recipient cells was subjected to 10 Gy IR. Cells were harvested for immunoblotting or fixed for immunofluorescence 2h or 6h post-IR treatment.

### Western blot

Cells were centrifuged at 4°C and 500g for 5 mins, before lysis in 5x the cell pellet volume of RIPA buffer (Thermo Scientific #89901), supplemented with 1x protease inhibitor (Merck # 11873580001) and 1x phosphatase inhibitor (Roche # 4906837001). Samples were sonicated for 10-30s and their concentration determined using the Pierce BCA Protein Assay Kit (Thermo Scientific #10678484), according to manufacturer’s protocol.

Cell lysates were diluted to equal concentrations with ddH_2_O and Laemmli buffer with β-mercaptoethanol (Alfa Aesar #J61337.AD) added to a final 1x concentration. Samples were denatured at 95°C for 5 mins and standard western blotting carried out using 4-15% Mini-PROTEAN TGX Precast protein gels (BioRad #4561083) and nitrocellulose membranes (Perkinelmer #NBA085A001EA). Total protein staining was carried out with Ponceau S (Merck #P7170-1L) for 5 mins. Bands were visualised using ECL substrate (Thermo Scientific #32106) and Amersham ECL Hyperfilm (Cytiva # 28-9068-35) according to manufacturer’s instructions. If applicable, the Super Signal Western Blot Enhancer kit (Thermo Fisher # 46640) was used according to the manufacturer’s instructions. Blots were quantified using ImageJ.

### Immunofluorescence

Following any necessary treatment, cells were fixed and permeabilised for 10 mins each at room temperature with 4% paraformaldehyde in PBS (Alfa Aesar #J61899) and 0.2% Triton X-100 (Sigma #X100-500ML) in PBS (0.2% PBST), respectively. Blocking was carried out using 10% FBS in PBS for 2h at 4°C. Primary antibodies were diluted in 10% FBS in PBS then incubated with coverslips overnight at 4°C in a humidified chamber. Washes were carried out in 0.1% PBST, before incubation with appropriate secondary antibodies diluted in 0.15% FBS in PBS for 2h at room temperature in a humidified chamber. Coverslips were mounted using Mounting Medium with DAPI (Abcam #ab104139-20ml). Slides were imaged using Olympus Fluoview FV1200 confocal microscope with a 60x objective lens, and a numerical aperture of 1:40. Images were quantified using CellProfiler 4.2.1 using a custom pipeline.

### Polymerase chain reaction

Genomic DNA was extracted using Phenol/chloroform/isoamyl alcohol (25:24:1, v/v) (Invitrogen #15593031) and precipitated with isopropanol. Semi-nested PCR was performed on genomic DNA to verify YBX1 knockout, as described by Shurtleff *et al.(Shurtleff et al., 2016)*

### RNA isolation

Total RNA was isolated from whole cell extracts, nuclear/cytoplasmic fractions or streptavidin beads following RNA immunoprecipitation. Samples were combined with an equal volume of TRIzol LS (Invitrogen # 10296028) followed by addition of 1-Bromo-3-chloropropane (Sigma-Aldrich # B9673) according to manufacturer’s instructions. Isopropanol precipitation was carried out by overnight incubation at-20°C, in the presence of 75mM NH_4_OAc and 2μl GlycoBlue. RNA pellet was washed sequentially with 100% ethanol, 70% ethanol and 100% ethanol, before resuspension in DEPC-treated water.

Small RNA from EV samples was also isolated by TRIzol LS extraction. This was followed by ethanol precipitation was using the mirVana miRNA Isolation Kit (Life Technologies #AM1561), following the manufacturer’s small RNA enrichment procedure. Elution was carried out twice, in 30μl DEPC-treated water each time.

### Small RNA sequencing

RNA was isolated from EV samples obtained from WT non-irradiated, WT irradiated and YBX1-/-irradiated cells, and quality controlled. Libraries were constructed using NEBNext® Multiplex Small RNA Library Prep Set (New England BioLabs) and sequenced using NextSeq 500.

### Northern blot

RNA samples were prepared to equal concentrations in 2x TBE-Urea Sample Buffer (Thermo Scientific # LC6876) and denatured at 98°C for 3 minutes. Separation was carried out on 12% denaturing (50% g/v urea) polyacrylamide gels in 1x TBE at 450V for 2.5h. RNA was transferred to positively charged nylon membranes (Amersham Hybond-N+ #RPN203B) at 5V for 1h in 0.5x TBE, followed by UV-crosslinking and pre-hybridisation in pre-warmed ULTRAhybOligo buffer (Thermo Fisher #AM8663) at 42°C for at least 1h. DNA probes against target RNA were radiolabelled with ^32^P-ATP (Perkinelmer #BLU502Z500UC), using T4-polynucleotide kinase (NEB #M0201L), for 2h at 37°C. Radiolabelled probes were purified using Microspin G-25 columns (Merck #GE27-5325-01) and added to membranes for overnight incubation at 42°C with rotation. 2x washes in 0.1% SSC buffer were then carried out for 15 mins each, before subjection to autoradiography with Amersham Hyperfilm (VWR #28-9068-45).

### siRNA, ASO, plasmid and synthetic oligonucleotide transfection

siRNA was reverse-transfected using Lipofectamine™ RNAiMax (Invitrogen #13778075) according to the manufacturer’s protocol, at a concentration of 60nM. Reverse transfection of ASOs was carried out in the same way, to a final concentration of 125nM. Plasmid reverse transfections were carried out using Lipofectamine™ LTX and Plus Reagent (Invitrogen #15338100) according to manufacturer’s instructions. Plasmids were transfected at a concentration of 500ng-1µg. Unmodified ysRNA oligonucleotides were forward transfected at a concentration of 10nM using Lipofectamine™ RNAiMax. AlexaFluor488-tagged ysRNA oligonucleotides were reverse transfected using Lipofectamine™ RNAiMax at concentrations of 10nM.

### RNA immunoprecipitation

Following necessary treatments, cells were pelleted by centrifugation at 500g and 4°C for 5 mins, then lysed in 5x pellet volume if RIP lysis buffer, supplemented with Ribolock RNase inhibitor (Life Technologies #EO0382) and DNase I (NEB #M0303S). Lysates were incubated at 4°C for 30 mins with rotation, then centrifuged at 17,000g and 4°C for 10 mins. Supernatant was kept and 10% removed for input. Alternatively, subcellular fractionation was carried out as described below.

Lysates were pre-cleared with streptavidin magnetic beads (NEB #S1420S) for 2h at 4°C, then incubated with 1μg relevant biotinylated RNA in 2x TENT buffer supplemented with Ribolock, 1x protease inhibitor and 1x phosphatase inhibitor for 30 mins at room temperature. Fresh streptavidin magnetic beads were then added to each sample and pulldown was carried out for 1.5h at 4°C with rotation. Where biotinylated RNA had been transfected into cells, streptavidin magnetic beads (NEB) were added directly to each lysate. Pulldown was incubated for 1h at 4°C with rotation. For analysis of immunoprecipitated proteins, beads were washed 4x in ice-cold 1x TENT buffer and eluted in 2x Laemmli buffer by incubation at 95°C for 10 mins. For analysis of immunoprecipitated RNA, beads were washed 3x in ice-cold 1x TENT buffer then 2x in ice-cold PBS. Beads were resuspended in 100μl PBS then eluted by incubation with Trizol LS reagent. RNA isolation was then carried as described previously.

### Subcellular fractionation

Cells were harvested from 15cm dishes by centrifugation at 500g and 4°C for 5 mins. All buffers were supplemented with 1x protease and phosphatase inhibitors, and 2μl/ml Ribolock RNase inhibitor. Cell pellet was lysed in 5x volume of hypotonic lysis buffer (10 mM HEPES pH 7.9, 60 mM KCl, 1.5 mM MgCl_2_, 1 mM EDTA, 1 mM DTT, 0.075% NP-40) and incubated at 4°C for 10 mins with rotation. Nuclei were pelleted by centrifugation at 800g and 4°C for 5 mins, and the supernatant collected as the cytoplasmic fraction. Insoluble nuclear remnants were removed by centrifugation at 13,000rpm and 4°C for 5 mins. The nuclear pellet was washed 5x in hypotonic lysis buffer without NP-40, lysed in 1x volume of nuclear lysis buffer (20 mM HEPES pH 7.9, 400 mM NaCl, 1.5 mM MgCl_2_, 0.2 mM EDTA, 1 mM DTT, 5% glycerol) supplemented with RNase inhibitor and diluted with 2x volume of dilution buffer (20 mM HEPES pH 7.9, 1.6% Triton-X-100, 0.2% sodium deoxycholate). Nuclear lysates were sonicated for 10s at 10 microns, then centrifuged for 10 mins at 13,500rpm and 4°C. The supernatant was collected as the soluble nuclear fraction.

### Slot blot

Immunoprecipitated RNA samples were diluted in TE buffer and denatured at 95°C for 3 mins. Positively charged nylon membranes (Amersham Hybond-N+ #RPN203B) were pre-equilibrated in TE buffer and assembled within Bio-Dot Microfiltration Apparatus (Biorad #1706542), as per manufacturer’s instructions. Each slot was washed once in TE buffer, then RNA applied to the membrane and a gentle vacuum applied. Slots were washed in TE buffer then left to dry for 10 mins. UV-crosslinking was carried out at 2000J, followed by blocking in 2% milk in PBS for 1h at room temperature with shaking. Primary antibodies were diluted in 2% milk in PBS and incubated with membrane overnight at 4°C, with shaking. This was followed by 1hr incubation with appropriate secondary antibodies diluted in 2% milk in PBS at room temperature with shaking. RNA was visualised by autoradiography using ECL substrate and Amersham ECL Hyperfilm according to manufacturer’s instructions. Total RNA staining was then carried out via methylene blue incubation for 5 mins at room temperature. Blots were quantified using ImageJ.

### Proximity ligation assay

Cells were treated as appropriate, then fixed and permeablised for 10 mins each at room temperature with 4% paraformaldehyde in PBS (Alfa Aesar #J61899) and 0.2% Triton X-100 (Sigma #X100-500ML) in PBS, respectively. Duolink Proximity Ligation Assay (Sigma Aldrich #DUO92101-1KT) was carried out on fixed cells according to manufacturer’s protocol. Coverslips were mounted using Duolink In Situ Mounting Medium with DAPI (Sigma-Aldrich #DUO82040), and stored protected from light at 4°C. Slides were imaged using Olympus Fluoview Spectral FV1200 confocal microscope with a 60x or 100x oil immersion objective lens, and a numerical aperture of 1:40. Quantification was predominantly carried out using CellProfiler and the Speckle Count pipeline. If background measurements were high, ImageJ Find Maxima function was used with a cut-off of 700.

### MTT assay

Transfected HeLa cells were reseeded into 96-well plates at a density of 10,000 cells per well, 24h post-transfection. 48h post-transfection, plates were subject to 5Gy irradiation, if applicable. MTT Cell Proliferation Assay Kit (Abcam #ab211091) was used according to manufacturer’s instructions, with 100% DMSO used instead of MTT solvent. Absorbance at 590nm was measured at 24, 48 and 72h post-IR.

### Clonogenic assay

HeLa cells were transfected with ASOs in 6-well plates, as described above. 48h after transfection, cells were treat with 2Gy IR and left to grow for 7-14 days until visible colonies formed. Plates were then washed twice in PBS, incubated with 0.5% crystal violet (Sigma #C6158-100G) in 20% ethanol for 30 mins, rinsed in water and left to dry. Plates were scanned and images quantified using the ColonyArea ImageJ plugin.

### Co-immunoprecipitation

Following transfection with YFP-PARP1 plasmid, cells from 15cm dishes were washed 3x in ice cold PBS, scraped in PBS and pelleted by centrifugation at 500g and 4°C for 5 mins. Cell pellets were lysed in 5x volume of co-IP lysis buffer, supplemented with Pierce Universal Nuclease. Lysates were incubated at 4°C for 30 mins with rotation, followed by centrifugation at 17,000g and 4°C for 10 mins. Supernatant was diluted in 1.5x volume co-IP dilution buffer and 10% removed as input. GFP-Trap beads were washed 3x and resuspended in cold co-IP dilution buffer. 50μl beads were added to each sample and incubated for 2h at 4°C with rotation. Supernatant was then removed and beads washed 3x in cold co-IP dilution buffer, followed by elution in 2x Laemmli buffer at 95°C for 10 mins.

### Bioinformatic analysis smallRNA-Seq Data Processing

Adapters were trimmed using Cutadapt (version 4.4) (https://cutadapt.readthedocs.io/en/stable/installation.html) in paired end mode and the quality of the resulting fastq files were assessed using FastQC (https://www.bioinformatics.babraham.ac.uk/projects/fastqc/). The trimmed reads were then aligned to GENCODE GRCh38 reference genome using STAR aligner(Dobin et al., 2013). Custom BED file containing GENCODE GRCh38 annotation with all RNA species except miRNAs were created using custom python script. Read counts across various smallRNA species (snoRNA,snRNA,tRNA, vaultRNA,YRNA) for each alignment file was calculated using BEDTools(Quinlan and Hall, 2010) multicov function. DESeq2(Love et al., 2014) was then used to perform differential expression analysis on the counts obtained using all three replicates for each condition.

### tRFs Analysis

MINTmap(Loher et al., 2017) was used to obtain read counts for each annotated tRNA-derived fragment (tRF) from MINTBase(Pliatsika et al., 2018). The read counts were then normalized to CPM based on total no of tRFs obtained for each sample. T test was then used to compare enrichment of various tRF species between the conditions (n=3) with significance threshold set at p value < 0.05 and |log2Fold Change| > 0.1. Differential enrichment of tRFs was then visualized using EnhancedVolcano (https://bioconductor.org/packages/devel/bioc/vignettes/EnhancedVolcano/inst/doc/EnhancedVolc ano.html) R package(www.R-project.org). Foldchange of commonly DE enriched tRFs between various conditions were plotted as a heatmap using Matplotlib python package

### miRNA Analysis

GRCh38 Co-ordinates for mature miRNAs annotation were obtained from miRBase(Kozomara and Griffiths-Jones, 2014). BEDTools multicov was then used to obtain read counts across mature miRNAs for each of the alignment files resulting from alignment of trimmed reads to GENCODE GRCh38. T test was then used to compare enrichment of various miRNA species between the conditions (n=3) with significance threshold set at p value < 0.05 and |log2Fold Change| > 0.1. Differential enrichment of miRNAs was then visualized using EnhancedVolcano (https://bioconductor.org/packages/devel/bioc/vignettes/EnhancedVolcano/inst/doc/EnhancedVolcano.html) R package. Foldchange of commonly DE enriched miRNAs between various conditions were plotted as a heatmap using Matplotlib python package.

### Metagene Plots

BigWig files containing RPKM normalized read count per nucleotide position was generated for each alignment file using deepTools bamCoverage(Ramirez et al., 2014). ComputeMatrix operation of deepTools was then performed on the bigWig files to calculate the RPKM coverage across annotated YRNA species. Matrices were then visualized using plotProfile function of deepTools.

### smallRNA Proportion plots

Raw coverage across mature miRNA, tRNAs, snoRNAs, snRNAs, YRNAs and vaultRNA were plotted as a pi-chart using Matplotlib python package.

### Fragment size distribution plots

BEDTools getfasta was used to extract all reads mapping to co-ordinates of various YRNA species. Custom python script was then used to plot percentage of fragments for all possible fragment sizes engendered from fragmentation of YRNA.

### Data and statistical analysis

All statistical analyses were performed in GraphPad Prism 9.3.1. All error bars represent mean ± SEM unless otherwise stated. Statistical testing was performed using the Student’s *t*-test, one-way ANOVA, Mann–Whitney test for two group non-parametric comparison (for PLA foci analysis), or Kruskall-Wallis test with Dunn’s correction for multiple group comparisons. Significances are listed as \**p* ≤ 0.05, \*\**p* ≤ 0.01, \*\*\**p* ≤ 0.001, \*\*\*\**p* ≤ 0.0001.

